# Diet prevents the expansion of segmented filamentous bacteria and ileo-colonic inflammation in a model of Crohn’s disease

**DOI:** 10.1101/2022.07.06.498810

**Authors:** Amira Metwaly, Jelena Jovic, Nadine Waldschmitt, Sevana Khaloian, Helena Heimes, Deborah Häcker, Nassim Hammoudi, Lionel Le Bourhis, Aida Mayorgas, Kolja Siebert, Marijana Basic, Tobias Schwerd, Matthieu Allez, Julian Panes, Azucena Salas, André Bleich, Sebastian Zeissig, Pamela Schnupf, Fabio Cominelli, Dirk Haller

**Affiliations:** Chair of Nutrition and Immunology, Technical University of Munich, Germany; APHP, Hôpital Saint Louis, Department of Gastroenterology, INSERM UMRS 1160, Paris Diderot, Sorbonne Paris-Cité University, France; Department of Experimental Pathology, Instituto de Investigaciones Biomédicas de Barcelona CSIC, IDIBAPS, CIBERehd, Spain; Department of Pediatrics, Dr. von Hauner Children’s Hospital, University Hospital, LMU Munich; Hannover Medical School, Institute for Laboratory Animal Science, Hannover, Germany; Department of Medicine I, University Hospital Dresden, Technische Universität (TU) Dresden, Dresden, Sachsen, Germany; Université Paris Cité, INSERM UMR-S1151, CNRS UMR-S8253, Institut Necker Enfants Malades, F-75015 Paris, France; Digestive Health Research Institute, Case Western Reserve University School of Medicine, Cleveland, OH, USA; ZIEL-Institute for Food and Health, Technical University of Munich, Germany

**Keywords:** Crohn’s disease, segmented filamentous bacteria, purified diet, *Tnf* ^ΔARE^ mice, inflammation, pathobiont, IBD

## Abstract

Crohn’s disease (CD) is associated with changes in the microbiota, and murine models of CD-like ileo-colonic inflammation depend on the presence of microbial triggers. Increased abundance of unknown Clostridiales and the microscopic detection of filamentous structures close to the epithelium of *Tnf* ^ΔARE^ mice pointed towards segmented filamentous bacteria (SFB), a commensal well-known to induce the maturation of Th17 cell-derived immune responses that is highly implicated in the pathogenesis of IBD. We show that the abundance of SFB strongly correlates with the severity of CD-like ileal inflammation in *Tnf* ^ΔARE^ and SAMP/Yit mice. SFB mono-colonization of germ-free *Tnf* ^ΔARE^ mice confirmed the causal link and resulted in severe ileo-colonic inflammation, characterized by elevated tissue levels of *Tnf* and *Il-17*, neutrophil infiltration and loss of Paneth and goblet cell function. Co-colonization of SFB in human-microbiota associated *Tnf* ^ΔARE^ mice confirmed that SFB presence is indispensable for disease development. Screening of 412 ileal and colonic mucosal biopsies from IBD patients using previously published and newly designed human SFB-specific primer sets showed no presence of SFB in human tissue samples. Simulating the protective effect of exclusive enteral nutrition (EEN) by feeding SFB mono-colonized *Tnf* ^ΔARE^ mice EEN-like purified diet antagonized SFB colonization and prevented disease development in *Tnf* ^ΔARE^ mice, clearly demonstrating the important role of diet in modulating this IBD-related but murine pathobiont.

## INTRODUCTION

CD is one of the two main subtypes of inflammatory bowel diseases (IBD) that affect the gastrointestinal tract and is characterized by patchy transmural inflammation along the small intestine, predominantly affecting the terminal ileum and proximal colon. Accumulating evidence proved that CD is a multifactorial disease, mainly driven by complex interactions between genetics, environmental factors, gut microbiota, and immune responses (Kaplan, 2015; Renz et al., 2011). To date, more than 70 CD and 241 IBD susceptibility loci have been identified via large-scale genome-wide association studies (GWAS) (Franke et al., 2010; Katrina M de Lange, 2017). Of note, many of these genetic risk loci are involved in immune pathways linked to microbial sensing and signaling, lack of adaptive mechanisms (e.g. interleukin (IL) 23/T helper (Th) 17 pathway) in the regulation of immune activation, autophagy, and Paneth cell function (Graham and Xavier, 2020; Jostins et al., 2012; Liu et al., 2015). However less than 30% of the heritability to CD is explained by known genetic variants, suggesting that microbial triggers play a dominant role in disease etiology. Perturbations of gut microbiome structure and function (also referred to as dysbiosis) are causally linked to a distortion of microbe-host homeostasis, leading to the development of chronic intestinal inflammation, such as Crohn’s disease (CD) and Ulcerative colitis (UC) (Ianiro and Hansen, 2018; Kamada et al., 2013; Levy et al., 2017; Metwaly et al., 2022). In this context, work from us and others clearly demonstrated that the presence of commensal bacteria is required to induce inflammation in several mouse models of IBD, while germ-free (GF) counterparts remained completely disease-free (Lengfelder et al., 2019; Metwaly et al., 2020; Rehaume et al., 2014; Roulis et al., 2016; Schaubeck et al., 2016).

The *Tnf* ^ΔARE^ CD mouse model is one of the very few models that develop intestinal inflammation resembling the transmural ileal disease phenotype frequently seen in CD patients (Ahmed et al., 2021; Buttó et al., 2015; Cominelli et al., 2017; Kontoyiannis et al., 1999). We previously showed that GF *Tnf* ^ΔARE^ mice are completely disease-free, and the transfer of disease-associated intestinal microbiota induced CD-like inflammation (Schaubeck et al., 2016). Further, mucosa-associated microbial communities of *Tnf* ^ΔARE^ mice were characterized by the expansion of segmented filamentous bacteria (SFB) (Roulis et al., 2016), a spore-forming, anaerobic symbiotic commensal bacteria of the phylum Firmicutes. SFB were shown to attach to the ileal mucosa at the time of weaning and are responsible for controlling immune cell maturation, including IL17-producing innate and adaptive lymphocytes (Al Nabhani et al., 2019; Gaboriau-Routhiau et al., 2009; Lécuyer et al., 2014; Schnupf et al., 2015). Although first reported over 100 years ago, SFB can only be cultured for a limited amount of time *in vitro* in an epithelial cell-SFB co-culturing system (Schnupf et al., 2015). Using 16S rRNA gene sequence analysis, it has been shown that SFB of mice, rats, and chickens represent a cluster within the *Clostridium* genus (Snel et al., 1998). First evidence for the presence of SFB in humans was based on the visualization of a tentative SFB organism adherent to ileal tissue biopsies by light microscopy (Klaasen et al., 1993) and this was later supported in samples from patients with ulcerative colitis (Caselli et al., 2013). More recently, 16S rRNA gene sequences of SFB were reported in human samples using SFB-specific PCR primers, where SFB sequences were found in 55 fecal samples (Yin et al., 2013) and in one ileostomy sample (Jonsson, 2013) . Further, Finotti et al. (2017) reported detection of SFB in tissue biopsies from the terminal ileum of UC patients (Finotti et al., 2017). In mice, SFB adhesion to intestinal epithelial cells contributes to the mucosal immune system function by inducing the differentiation of Th17, production of secretory immunoglobulin A (IgA), and the secretion of antimicrobial peptides (AMPs) (Atarashi et al., 2015; Ivanov et al., 2009; Salzman, 2010). Similar associations were observed in luminal fluids from children, suggesting comparable immunomodulatory effects of SFB growth in early age of human individuals (Chen et al., 2018). Although the role of SFB in humans to induce antigen specific Th17 cells remains largely unknown, they appear to play an important role in modulating the gut immune response in mouse models of ileal inflammation. The regulatory mechanisms controlling SFB growth remain unclear.

Diet is proposed to be an essential factor in IBD that targets inflammation-related mechanisms in the host, either directly or indirectly via changes in the structure and function of the intestinal microbiota (Khalili et al., 2018; Schaubeck and Haller, 2015).Studies on various patient cohorts proposed specific dietary components to be associated with IBD, including higher consumption of sugar and animal fat as well as a reduced vegetable intake (Ananthakrishnan et al., 2014; Racine et al., 2016) . Early studies in the 1980’s suggested that dietary composition might be an important determinant of the presence of SFB in the small intestine of mice, however the dietary components responsible were not identified (Klaasen et al., 1991; Koopman et al., 1987, 1986). Here, we identified a novel pathogenic role of SFB in driving severe CD-like ileo-colonic inflammation characterized by loss of Paneth and goblet cell functions in *Tnf* ^ΔARE^ mice. A purified diet antagonized SFB colonization and prevented disease development in *Tnf* ^ΔARE^ mice in contrast to a fiber-containing chow diet, clearly demonstrating the important role of diet in modulating a novel IBD-relevant pathobiont. Taken together, we showed that dietary intervention using EEN-like purified diet in a murine model of CD prevented pathobiont-mediated intestinal inflammation, supporting a direct link between diet and microbial communities in mediating protective functions.

## METHODS

### Ethics statement

Mouse experiments and the treatment protocols were approved by the Committee on Animal Health and Care of the local government body of the state of Upper Bavaria (Regierung von Oberbayern; approvals number (55.2-1-54-2531-99-13, 55.2-1-54-2532-133-2014 and 55.2-2532.Vet_02-17-104) and performed in compliance with the EEC recommendations for the care and use of Lab. Animals. (European Communities Council Directive of 24 November 1986 (86/609/EEC). Animals were housed in the germ-free (GF) mouse facility at the Technical University of Munich (School of Life Sciences). IBD Patients were recruited at the Department of Gastroenterology, Hospital Clinic Barcelona, at Hospital San Louis in Paris and at Dr. von Hauner Children’s Hospital in Munich. The study was approved by the corresponding hospital ethics committee. All patients provided written informed consent.

### Animal experiments

*Tnf* ^ΔARE^ mice and wild-type (WT) control littermates were imported from Case Western Reserve University in Cleveland and housed under germ-free (GF) conditions. 8 weeks old GF mice were colonized with specific-pathogen-free (SPF)-derived WT cecal microbiota and maintained under SPF housing conditions. Co-housing experiments were performed with 8 weeks old female *Tnf* ^ΔARE^ and WT littermates and female SFB mono-colonized NOD-SCID mice. Monocolonization of ARE and WT mice with SFB was performed either by breeding of mice with SFB-mono-colonized WT mothers or by oral gavage of gut content from mono-colonized NOD-SCID mice into 8 weeks old *Tnf* ^ΔARE^ mice and controls. Mono-colonization of mice with *Alistipes*, *Lactobacillus murinus*, CD patient derived *E. coli* LF82 was done by oral gavage of cultured bacteria. Humanization experiment was performed by transferring human fecal microbiota to *Tnf* ^ΔARE^ and WT mice at the age of 8 weeks. Preparation of fecal material for inoculation in germ-free mice was done under anaerobic conditions and using reduced PBS (PBS supplemented with 0.05% L-cysteine-HCl) in an anaerobic chamber (atmosphere, 90% N2, 5% CO2, and 5% H2) and vortexed at room temperature for 5 minutes. The fecal suspension was allowed to settle by gravity for 5 minutes to exclude residual particulate matter. The clear supernatant was transferred under anaerobic conditions into an anaerobic crimped tube, which was transferred to the gnotobiotic facility. Each germ-free recipient mouse received 100 µL of the suspension via oral gavage using a 20 Gauge gavage needle (Fine Science Tools). Mice were sacrificed 4 weeks after colonization. WT and *Tnf* ^ΔARE^ mice were placed on purified diets (Ssniff E15000, Germany), enriched with fiber (Ssniff S5745-E907, Germany) at 7 or 9 weeks of age. SFB mono-colonization was performed by transplantation of gut content of mono-colonized NOD-SCID mice at 8 weeks of age, and mice were sampled at 12 weeks of age. Detailed information about diet ingredients is summarized in **Supplementary Table S1 .**

Germ-free WT and IL-10-deficient (*Il10*^−/−^) mice on 129Sv/Ev background were kept at the gnotobiology core facility of the Institute for food & health, Technical University Munich, Germany. Mono-colonization of *Il10*^−/−^ and WT mice with SFB was performed by oral gavage of gut content from mono-colonized NOD-SCID mice into 8 weeks old *Il10^−/−^* mice and WT controls.

SAMP1Yi/Fc (SAMP) mice: Ten- and >24-week-old specific-pathogen-free (SPF) SAMP and age/sex matched AKR/J mice were used. Mice were kept on a 12-h:12-h light: dark cycle in a species-appropriate temperature/humidity-controlled room and maintained in AAALAC-accredited Animal Research Center rooms at Case Western Reserve University (CWRU).

### Housing of germ-free mice

GF mice were kept at the gnotobiology core facility of the Institute for food & health, Technical University Munich, Germany. GF mice were housed in flexible film isolators ventilated via HEPA-filtered air at 22 ± 1°C with a 12-h light/dark cycle. Before experiments, littermates were combined and randomly assigned to treatment groups. A maximum of 5 mice are housed per cage (floor area ∼540 cm^2^). Mice received standard Chow, purified or fiber-rich purified diet (autoclaved V1124-300; Ssniff, Soest, Germany) and autoclaved water *ad libitum*.

### CD Patients and control subjects

Mucosal biopsies for SFB quantification were collected from three IBD patient cohorts enrolled in Barcelona, Paris and Munich **Supplementary Table S2**. ***Spanish (HSCT) cohort***: enrolled patients were in follow-up for CD following treatment with autologous hematopoietic stem cell transplantation (AHSCT). Ileal (n=42) and colonic biopsies (n=84) from CD patients were included in the analysis. As a control, ileal and colonic biopsies (n=11 and 30 respectively) from a group of healthy subjects performing colonoscopy for screening purposes were included in the analysis. CD patients recruited fulfilled the previously described inclusion criteria (Corraliza et al., 2019; Jauregui-Amezaga et al., 2016; López-García et al., 2017; Metwaly et al., 2020). Biopsies were immediately frozen at −80°C for PCR analysis). ***French (Biotherapy) cohort***: This longitudinal study comprised patients with IBD (CD and UC) who are treated with anti-TNF and other biotherapy drugs and recruited in Paris (n=172), Ileal (n=40) and colonic biopsies (n=138) from CD and UC patients were included in the analysis. ***German pediatric cohort***: This longitudinal prospective study comprised children with IBD and were recruited in Munich (n=67), Ileal (n==32) and colonic biopsies (n=35) were included in the analysis.

### Bacterial cultivation and inoculation of mice

*Lactobacillus murinus, E. coli LF8*2 (Adherent Invasive *Escherichia coli* derived from a chronic ileal lesion in a CD patient) and *Alistipes sp*. JC136 were cultivated in Wilkins-Chalgren anaerobe broth (WCA, Oxoid) that was prepared in Hungate tube flushed with Nitrogen. Purity of bacterial strains was controlled by microscopic evaluation following a standard Gram staining procedure. GF age matched ARE and WT mice were inoculated with 1x 10^8 CFU by oral gavage and sacrificed 4 weeks after colonization. MIBAC – Minimal mouse Bacteriome is a consortium of 7 bacteria acts as a proxy model of a complex mouse gut microbiota (unpublished). It was selected based on prevalence and abundance values in mouse gut using omics datasets. The final composition of MIBAC is as follows: *Clostridium ramosum, Paraclostridium bifermentans, Enterococcus hirae, Enterorhabdus mucosicola, Escherichia coli, Lactobacillus murinus and Parabacteroides goldsteinii*.

### Bacterial DNA extraction from feces

Fecal pellets were resuspended in 600 µl DNA stabilization solution (STRATEC biomedical), 400 µl Phenol:Chloroform:IsoAmyl alcohol (25:24:1; Sigma Aldrich), and 500 mg autoclaved zirconia/silica beads (0.1mm; Roth). Bacterial cells were mechanically disrupted by using a FastPrep®-24 (MP Biomedicals) followed by a heat treatment for 8 min at 95 °C and centrifugation with 15000 x g for 5 min at 4 °C. Supernatants were treated with RNase (0.1 µg/ µl; VWR International) for 30 min at 37 °C. DNA was purified by using the gDNA clean-up kit (Macherey-Nagel). DNA concentrations and purity were checked using NanoDrop® (Thermo Fisher Scientific), and DNA integrity was checked by gel electerophoresis (0.5%, 100V, 45 min). Samples were immediately used or stored at −20 C for long-term storage.

### DNA extraction from tissue biopsy

DNA was extracted from biopsies by using the NucleoSpin® Tissue kit (Machery-Nagel). Briefly, tissue biopsies were resuspended in 180 µl sterile filtered lysis buffer (20 mM Tris/HCl; 2 mM EDTA; 1 % Triton X-100 (pH 8) and supplemented with freshly prepared lysozyme solution (20 mg/ml). Tissue suspensions were incubated in a thermoshaker (37°C, 30 min, 950 rpm). Afterwards, Proteinase K (10mg/ml) was added, and the suspension was vortexed vigorously and incubated in a thermoshaker (56°C, 1-3 h, 950 rpm) until complete lysis of tissue pieces. Further purification steps were performed by using the NucleoSpin® Tissue kit following the manufacturer’s instructions.

### High throughput 16S ribosomal RNA (rRNA) gene sequencing and microbiome profiling

Library preparation and sequencing were performed as described in detail previously. In brief, the V3-V4 regions of the 16S rRNA gene was amplified (10×15 cycles for fecal samples, 15×15 cycles for tissue biopsies) by using previously described two-step protocol (Berry et al., 2011) using forward and reverse primers 341F-785R 34. Purification of amplicons was performed by using the AMPure XP system (Beckmann). Sequencing was performed with pooled samples in paired-end modus (PE275) using a MiSeq system (Illumina, Inc.) according to the manufacturer’s instructions and 25 % (v/v) PhiX standard library. Processing of raw reads was performed by using IMNGS pipeline based on the UPARSE approach (Edgar, 2013). Sequences were demultiplexed, trimmed to the first base with a quality score <3 and then paired. Sequences with less than 300 and more than 600 nucleotides and paired reads with an expected error >3 were excluded from the analysis. Trimming of remaining reads was done by trimming 5 nucleotides on each end to avoid GC bias and non-random base composition. A table of zOTUs is constructed by considering all reads before any quality filtering. ZOTUs (zero-radius OTUs) are valid operational taxonomic units that provide the maximum possible biological resolution compared to conventional 97% OTUs. Since using 97% identity may merge phenotypically different strains with distinct sequences into a single cluster, zOTUs are found to be superior to conventional OTU clusters ((Edgar, 2016)Our) . Taxonomy assignment was performed at 80 % confidence level using the SILVA ribosomal RNA gene database project (Yilmaz et al., 2014). Analysis was performed in R programming environment by using Rhea R-package (Lagkouvardos et al., 2017). Sequencing depth was assessed by using rarefaction curves, and samples with low quality were removed or re-sequenced. Diversity between groups was analyze by using ß-diversity based on generalized UniFrac distances. Alpha-diversity was assessed based on species richness and Shannon effective diversity. P-values were calculated by using ANOVA on Ranks and corrected for multiple comparisons according to the Benjamini-Hochberg method. Only taxa with a prevalence of at least 30 % samples in one given group were considered for statistical analysis.

### Reverse transcriptase-polymerase chain reaction

Total RNA from total ileal, caecal and colonic tissue was isolated by using the NucleoSpin RNAII kit (Macherey-Nagel GmbH) according to manufacturer’s instructions. Complementary DNA was synthetized from 500 ng total RNA by using random hexamers and moloney murine leukemia virus (M-MLV) reverse transcriptase (RT) Point Mutant Synthesis System (Promega). Quantification was performed by using the LightCycler 480 Universal Probe Library System (Roche). Following primer sequences and respective probes () were used: *Tnf* for_*5’-TGCCTATGTCTCAGCCTCTTC-3’, rev_5’-GAGGCCATTTGGGAACTTCT-3’* (49), *Gapdh* for_*5’-CACACCCATCACAAACATGG-3’, rev_5’-GCCAAAAGGGTCATCATCTC-3’* (29), *Il-17* for_*5’-AGGGATATCTATCAGGGTCTTCATT-3’, rev_5’-TGTGAAGGTCAACCTCAAAGTC-3’* (50), *Ifng*for_*5’-AGCGTTCATTGTCTCAGAGCTA-3’, rev_5’-CCTTTGGACCCTCTGACTTG-3’* (63), *Ang4* for_*5’-CGTAGGAATTTTTCGTACCTTTCA-3’, rev_5’-CCCCAGTTGGAGGAAAGC-3’* (106); *Defa-5* for_*5’-CAGAGCCGATGGTTGTCAT-3’, rev_5’-TTTTGGGACCTGCAGAAATC-3’*(84), *Reg3b* for_*5’-TCATCACGTCATTGTTACTCCA-3’, rev_5’-TGGATTGGGCTCCATGAC-3’* (10), *Muc2* for_*5’-GGTCTGGCAGTCCTCGAA-3’, rev_5’-GGCAGTACAAGAACCGGAGT-3’* (66). Quantification of SFB in fecal mouse content was performed using the MasterMix SensiFAST™ SYBR (BIOLINE) and specific primer pair for_*5‘-GACGCTGAGGCATGAGAGCAT-3‘, rev_5‘-GACGGCACGGATTGTTATTCA*-3‘ (Prakash et al., 2011). Quantification of SFB in human tissue biopsies was performed using the MasterMix SensiFAST™ SYBR (BIOLINE) and 3 different primer sets: *for*_*5‘-TGTGGGTTGTGAATAGCAAT-3‘*(Finotti et al., 2017)*, rev_5‘-GCGAGCTTCCCTCATTACAAGG*-3’, *for_5‘-AGGAGGAGTCTGCGGCACATTAGC*-*3‘* (Shukla et al., 2015; Suzuki et al., 2004) *,rev_5‘* TCCCCACTGCTGCCTCCCGTAG-*3‘* (Jonsson et al., 2020; Snel et al., 1995) *and for,_ 5’-TGTAGGTTGTGAAGAACAAT-3’, rev_ 5’-GCGAGCTTCCCTCATTACAAGG-3’* (modified primer sequences, unpublished data, Pamela Schnupf).

### Gel electrophoresis

Agarose gel 1 v/w % containing 0.001% Gelred (InvivoGen) was prepared by using Tris Acetate-EDTA buffer. qPCR amplicons were separated at 100 Volt for 1 hour following DNA band visualization under UV light.

### Isolation of immune cell population from lamina propria

Lamina propria-resident immune cells from small intestine were isolated by digesting intestinal tissue with Collagenase VIII (Sigma-Aldrich). Shortly, Payer’s patches were removed, and epithelial layer was dissociated by incubating tissue pieces twice in 2mM EDTA/PBS for 20 min at 37°C with shaking (180 rpm). Afterwards, single cell suspension was prepared by digesting small intestinal tissue in complete RPMI medium (10% FCS, 1% P/S, 1% Glu) containing Collagenase VIII (0.6 mg/ml) for approximately 15 min at 37°C with shaking (180 rpm). Lamina propria-resident immune cells from large intestine were isolated by digesting intestinal tissue with Collagenase D (Roche), Collagenase V (Sigma-Aldrich), Dispase I (Gibco), and DNase I (Roche). Here, epithelial cells were removed by incubating tissue pieces twiese in 2mM EDTA/PBS for 15 min at 37°C with shaking (180 rpm). Next, single cell suspension was prepared by digesting large intestinal tissue in complete RPMI medium (10% FCS, 1% P/S, 1% Glu) containing Collagenase D (1,25 mg/ml), Collagenase V (0,85 mg/ml), Dispase I (1 mg/ml), and DNase (30 U/ml) for a maximum of 45 min at 37°C with shaking (180 rpm). Cell number of single cell suspensions (2% FCS/PBS) were assessed, and cells were kept at 4°C for immediate flow cytometry analysis.

### Flow cytometry analysis

Cells were stained and subsequently analysed by using a LSRII system (BD Biosciences). Dead cells were excluded with the Zombie GreenTM or Zombie NIRTM Fixable Viability Kit (BioLegend). Allophycocyanin-Cy7-conjugated anti-CD3 (17A2), PE-Cy7-conjugated anti-CD4 (RM4-5), PE-Cy7-conjugated anti-CD11b (M1/70), PE-conjugated anti-CD45 (30-F11), PerCP/Cyanine5.5-conjugated anti-mouse I-A/I-E (M5/114.15.2), Allophycocyanin-Cy7-conjugated anti-Ly6C (HK1.4), Allophycocyanin-conjugated anti-Ly6G (1A8), and PerCP/Cyanine5.5-conjugated anti-CD3 (17A2) were from BioLegend, Allophycocyanin-conjugated anti-CD25 (PC61.5), PE-conjugated anti-Foxp3 (FJK-16s), PE-Cy7-conjugated anti-IL17 (eBio17B7), and Allophycocyanin-conjugated anti-IFN gamma (XMG1.2) were from Thermo Fisher Scientific, and FITC-conjugated anti-CD4 (RM4-5) was from BD Biosciences. FcR block was done by using FcR blocking reagent, mouse (Miltenyi Biotec) and intracellular staining was performed by using eBioscience™ Foxp3 / Transcription Factor Staining Buffer Set (Thermo Fisher Scientific).

### Tissue fixation and staining procedures

Intestinal specimens were fixed in 4% formaldehyde/PBS for 24 h at RT, subsequently dehydrated (Leica TP1020), and embedded in paraffin (McCormick; Leica EG1150C). 5 µm-thick tissue sections were prepared, deparaffinized and Hematoxylin and Eosin (H&E) staining was performed by using a Leica ST5020 Multistainer system. For periodic acid Schiff (PAS) and Alcian blue (AB) staining, formalin-fixed paraffin-embedded (FFPE) tissue sections were deparaffinized and rehydrated. Alcian blue solution for acidic mucins (1% volume/volume in 3% acetic acid, pH 2.5) was applied for 15 min. Tissue sections were incubated with periodic acid solution (0.5% volume/volume) for 5 min and co-staining with Schiff’s reagent (Sigma-Aldrich) for neutral mucins was performed for 10 min. Nuclei were counterstained with hematoxylin for 1 min. Numbers of goblet cells (GC) were calculated as a total number per area of interest. Images were acquired by using the Digital microscope M8 and MicroPoint software (PreciPoint GmbH). For immunofluorescent (IF) staining, 3.5 µm-thick tissue sections were placed on SuperFrost PlusTM slides (Thermo Fisher Scientific), deparaffinized (Leica ST5020), and rehydrated. Antigen retrieval was performed in 10 mM sodium citrate buffer (pH 6.0) by steaming sections in a microwave oven (900 Watt) for 23 min. Slides were rinsed with H2O and sections were blocked for 1h with blocking buffer containing the serum (5%). Primary antibodies against E-Cadherin (ab76055, 1:300, Abcam) and Lysozyme (A0099, 1:1000, DAKO, Agilent) were directly applied and incubated overnight at 4°C. Fluorescently labelled secondary antibodies anti-rabbit (A10040, A31571, 1:200, Life Technologies) and anti-mouse (A11001, 1:200, Life Technologies) were incubated light-protected for 1 h at RT. DAPI (Sigma Aldrich) was added for nuclear staining. Tissue sections were mounted (BIOZOL Diagnostica) and analyzed by using the Flouview FV10i microscope (Olympus).

### Fluorescent in situ hybridization

Dissected, intact ileal and colonic tubes were fixed in Carnoy solution (60% dry MeOH, 30% dry chloroform, 10% acetic acid) overnight at RT. Dehydration of samples was performed by washes in dry MeOH, 100% EtOH, xylene/100% EtOH (1:1), and xylene. Dehydrated tissue was submerged in melted paraffin for 20 minutes and embedded. 9 µm-thick tissue sections were deparaffinized, rehydrated, and fixed in 10% formalin before permeabilizing in a lysozyme solution (40 mg/mL lysozyme in a filter-sterilized 20 mmol/L Tris/2 mmol/L EDTA/1.2 % volume/volume Triton-X100 buffer) for 45 minutes at 37°C. Tissue sections were incubated with Cy5-conjugated bacterial probe EUB338 (5′-GCTGCCTCCCGTAGGAGT-3′) overnight at 46°C. Sections were co-stained with anti-MUC2 (H-300, 1:100, Santa Cruz Biotechnology) for 1h at RT following 30 min of incubation with secondary antibody (1:200, A11034, Life Technologies) and nuclei were visualized by using DAPI (Sigma Aldrich).

### Histological scoring system

Scoring of H&E stained tissue sections was performed blindly by evaluating lamina propria mononuclear cell infiltration, crypt hyperplasia, goblet cell depletion, and architectural distortion (Erben et al., 2014; Katakura et al., 2005). Grade of inflammation is reflected by total scores from 0 to 12.

## RESULTS

### Severity of CD-like inflammation correlates with SFB abundance in *Tnf* ^ΔARE^ and SAMP/YitFc mice

Our previous data showed that GF *Tnf* ^ΔARE^ mice are completely disease-free, and only upon the transfer of complex microbial communities derived from inflamed *Tnf* ^ΔARE^ mice, they develop intestinal inflammation. Increased abundance of unknown Clostridiales (Schaubeck et al., 2016) and the microscopic detection of rod-like structures in close proximity to the epithelium of *Tnf* ^ΔARE^ mice (Roulis et al., 2016) pointed towards the relevance of SFB. A colony of non-inflamed GF *Tnf* ^ΔARE^ mice and littermate WT controls (F0) was colonized with cecal content of SPF-housed WT mice having SFB counts reduced to undetectable levels. Mice were subsequently transferred to SPF housing for breeding (F1, F2) and monitoring of inflammation. At 18 weeks of age, the newly introduced *Tnf* ^ΔARE^ mice gradually developed ileal inflammation. The number of responder mice, however, significantly increased over the following two breeding generations. Similarly, subsequent generations of SPF-housed *Tnf* ^ΔARE^ mice exhibited varying severity of ileal inflammation independent of cage and litter effect **(Suppl. Figure 1A).** As we previously reported, three inflammatory phenotypes upon bacterial colonization could be observed in *Tnf* ^ΔARE^ mice, including severe inflammation (here referred to as responders, R, inflammation score >4), mild inflammation (here referred to as low responders, LR, inflammation score <4), as well as no inflammation (here referred to as non-responders, NR, inflammation score 0) **(Suppl. Figure 1B, C)**. We investigated the progression of ileal inflammation as well as SFB abundance over time. Our data show that shortly after weaning all 3-4 weeks old *Tnf* ^ΔARE^ mice were disease-free and started to show mild tissue pathology at the age of 8 weeks **(Figure 1A).** With regard to the SFB growth, both 3-4 weeks old *Tnf* ^ΔARE^ mice and WT controls revealed high SFB titers in cecal content, which subsequently declined below detectable levels in aging littermate WT controls, while it only has continuously decreased in non-inflamed NR *Tnf* ^ΔARE^ mice **(Figure 1B).** Histopathological evaluation showed tissue pathology to be restricted to the ileal compartment **(Figure 1C).** To visualize bacterial structures in ileal tissue sections of non-inflamed (NR) and inflamed (R) *Tnf* ^ΔARE^ mice (F1), we performed fluorescence in situ hybridization (FISH) by staining ileum sections with bacteria domain-specific (EUB338) probes. Microscopic examination showed detection of filamentous bacteria close to the intestinal epithelium of responder mice **(Figure 1D)**. The inflammation in ileal tissue of responder mice is characterized by significantly enhanced *Tnf* and *Il17* transcript levels compared to non-inflamed WT controls **(Figure 1E).** A correlation analysis on the full cohort of 18 weeks old *Tnf* ^ΔARE^ mice (NR, LR, R) resulted in significant association between SFB abundance and severity of ileal tissue pathology (p<0.0001, r=0.671) **(Figure 1F).** Next to *Tnf* ^ΔARE^ mice, we quantified SFB abundances in SAMP1/YitFc mice, a unique mouse model that spontaneously develop CD-like “cobblestone” ileitis with 100% penetrance and within a well-defined time course (pre-ileitis, disease induction, and chronic ileitis) (Pizarro et al., 2011). Similarly, a correlation analysis on a cohort of SAMP1/YitFc mice showed significant association between SFB abundance and severity of ileal tissue pathology (p<0.01, r=0.5895) **(Figure 1G)**, demonstrating a potential relevance of SFB as an important specific pathogenic factor for ileal inflammation.

**Figure 1.**
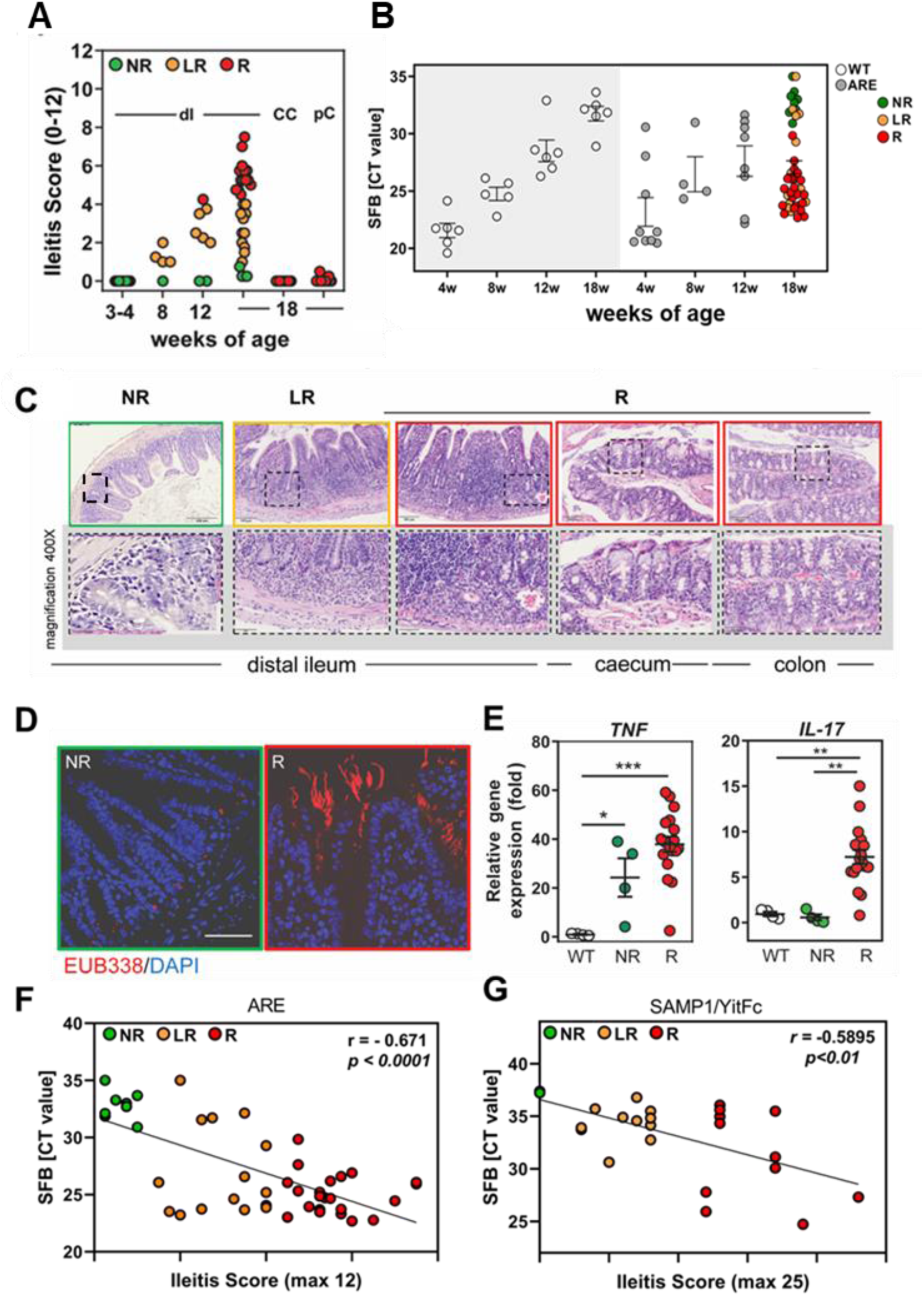
SFB abundance correlates with ileitis severity in SPF-house *Tnf* ^ΔARE^ mice. **(A)** Ileitis scores of SPF-housed *Tnf* ^ΔARE^ mice at the age of 3-4, 8, 12 and 18 weeks of age. Color-code represents severity of inflammation as described above. Cecal content of SPF-housed WT mice with undetectable SFB counts was introduced into germ-free *Tnf* ^ΔARE^ mice and littermate WT controls. **(B)** Quantitative Analysis of SFB abundance in *Tnf* ^ΔARE^ and matching WT mice overtime. CT value >30 is regarded as non-specificity threshold. **(C)** Representative H&E-stained tissue sections from distal ileum, caecum, and proximal colon of *Tnf* ^ΔARE^ mice showing no (NR), low (LR), and high grade (R) of inflammation in the ileum as described above. **(D)** Analysis of bacterial localization in distal ileum of inflamed (R) and non-inflamed (NR) *Tnf* ^ΔARE^ mice from F1 generation by performing FISH staining with eubacterial 16S probe (red) and DAPI for nuclear visualization. **(E)** Quantitative analysis of *Tnf* and *Il-17* mRNA expression in distal ileum of non-inflamed (NR) and inflamed (R *Tnf* ^ΔARE^ mice and WT controls. **(F)** Correlation analysis between Ileitis score and SFB abundance regarding the whole cohort of 18 weeks old *Tnf* ^ΔARE^ mice and **(G)** in twelve 10-week-old (mild ileitis) and twelve >24-week-old (severe ileitis) SAMP1/YitFc mice. Statistical significance was assessed by One-way-ANOVA followed by Tukey Multiple Comparison test. *p<0.05; **p<0.01; ***p<0.001 was considered statistically significant.

To examine SFB-driven alterations in complex microbial ecosystems, 8 weeks old, SPF-housed *Tnf* ^ΔARE^ mice and WT controls with low SFB counts in fecal content were co-housed with SFB-mono-colonized NOD-SCID mice for 10 weeks. Fecal pellets were collected weekly and analyzed using 16S rRNA gene amplicon sequencing. After 10 weeks of co-housing, all *Tnf* ^ΔARE^ mice showed significant signs of inflammation, while WT mice remained disease-free. To assess how SFB affects the microbiota, we compared gut microbiota composition between WT and *Tnf* ^ΔARE^ mice **(Suppl. Figure 1D)**. 16S microbial profiling revealed a substantial change in gut microbiota composition, characterized by significant increase in the relative abundance of *Alistipes* species in the inflamed *Tnf* ^ΔARE^ mice compared to WT controls. In contrast, increased abundance of *Lactobacillus* was evident for WT mice **(Suppl. Figure 1E)**. Intriguingly, subsequent mono-colonization of GF *Tnf* ^ΔARE^ mice with *Alistipes* failed to induce any inflammatory responses and/or ileal pathology **(Suppl. Figure 1F).** We additionally tested the impact of monocolonizing *Tnf* ^ΔARE^ mice with the human adherent-invasive *Escherichia coli* LF82 or *Lactobacillus spp*. and both showed no impact of these bacteria in inducing inflammation in *Tnf* ^ΔARE^ mice. To test the pathogenic potential of a more complex bacterial community, we colonized *Tnf* ^ΔARE^ mice and WT controls with a simplified human microbiota consortium, which comprises 7 bacterial strains (*Clostridium ramosum, Paraclostridium bifermentans, Enterococcus hirae, Enterorhabdus mucosicola, Escherichia coli, Lactobacillus murinus and Parabacteroides goldsteinii*) **(Supplmentary Table S3)** isolated from the mouse gut. However, and despite of the increased bacterial complexity, this bacterial consortium failed to induce inflammation in *Tnf* ^ΔARE^ mice.

### SFB promote intestinal inflammation in *Tnf* ^ΔARE^ mice

Next, we sought to investigate SFB-mediated immune modulation following high SFB exposure from birth. Therefore, GF WT mothers were monocolonized with SFB and mated with GF *Tnf* ^ΔARE^ males. Following maternal SFB exposure, mono-colonized offspring was analyzed at 4 and 12 weeks of age. Interestingly, 12 weeks old SFB-mono-colonized *Tnf* ^ΔARE^ mice showed extensive ileo-colonic inflammation affecting small and large intestinal compartment, whereas 4 weeks old mice remained completely disease-free **(Figure 2A).** Comparable SFB abundance was detected in all intestinal compartments of mono-colonized *Tnf* ^ΔARE^ and WT mice **(Figure 2B).** SFB was localized in close contact to the intestinal epithelium of the inflamed tissue regions, such as the distal ileum and proximal colon but did not penetrate the mucus layer of the distal colon **(Figure 2C).** Histopathological evaluation showed exacerbated inflammation in the ileum, cecum, and proximal colon, and no inflammation in the duodenum, jejunum, and distal colon **(Figure 2D).** SFB are well known to specifically induce Th17 responses in mice under physiological conditions (Ivanov et al., 2009). In SFB mono-colonized mice, analysis of the inflammatory transcript levels revealed highly enhanced expression of *Tnf*, *Il-17,* and *Ifng* in most of the affected mucosal tissue of 12 weeks old *Tnf* ^ΔARE^ mice when compared to WT controls, while *Tnf* was only found to be significantly elevated in the distal ileum of 4 weeks old *Tnf* ^ΔARE^ mice **(Figure 2E and Sup Suppl. Figure 2A).** Tissue pathology of inflamed *Tnf* ^ΔARE^ mice is characterized by a significantly increased immune cell infiltration, showing an increased number of IL-17-expressing cells in the tissue when compared to WT controls (**Suppl. Figure 2B**). However, the proportion of *Il-17*-expressing T cells remained unchanged between *Tnf* ^ΔARE^ mice and WT controls upon SFB colonization (**Suppl. Figure 2C)**. In contrast, considerable changes in cell proportions were observed for *Ifng*-expressing T cells, but also neutrophilic granulocytes, which were both found to be highly enhanced in 12 weeks old *Tnf* ^ΔARE^ mice (**Suppl. Figure 2D, E, F)**. Next, we determined the expression of genes involved in antimicrobial defense at the epithelium in the distal ileum to investigate the signaling molecules that are produced by the host upon SFB colonization in 4 weeks and 12 weeks old *Tnf* ^ΔARE^ mice. Twelve-week-old SFB mono-colonized *Tnf* ^ΔARE^ mice showed significantly reduced levels of *Muc2*, *Defa5*, *Ang4*, and *Reg3b* compared to 4-week-old mice or WT controls **(Figure 2F).** Consistent with these findings, significantly reduced numbers of lysozyme-positive Paneth cells, as well as mucin-filled goblet cells were shown in inflamed 12 weeks old *Tnf* ^ΔARE^ mice (**Figure 2G, H and Suppl. Figure 2H, I).**

**Figure 2.**
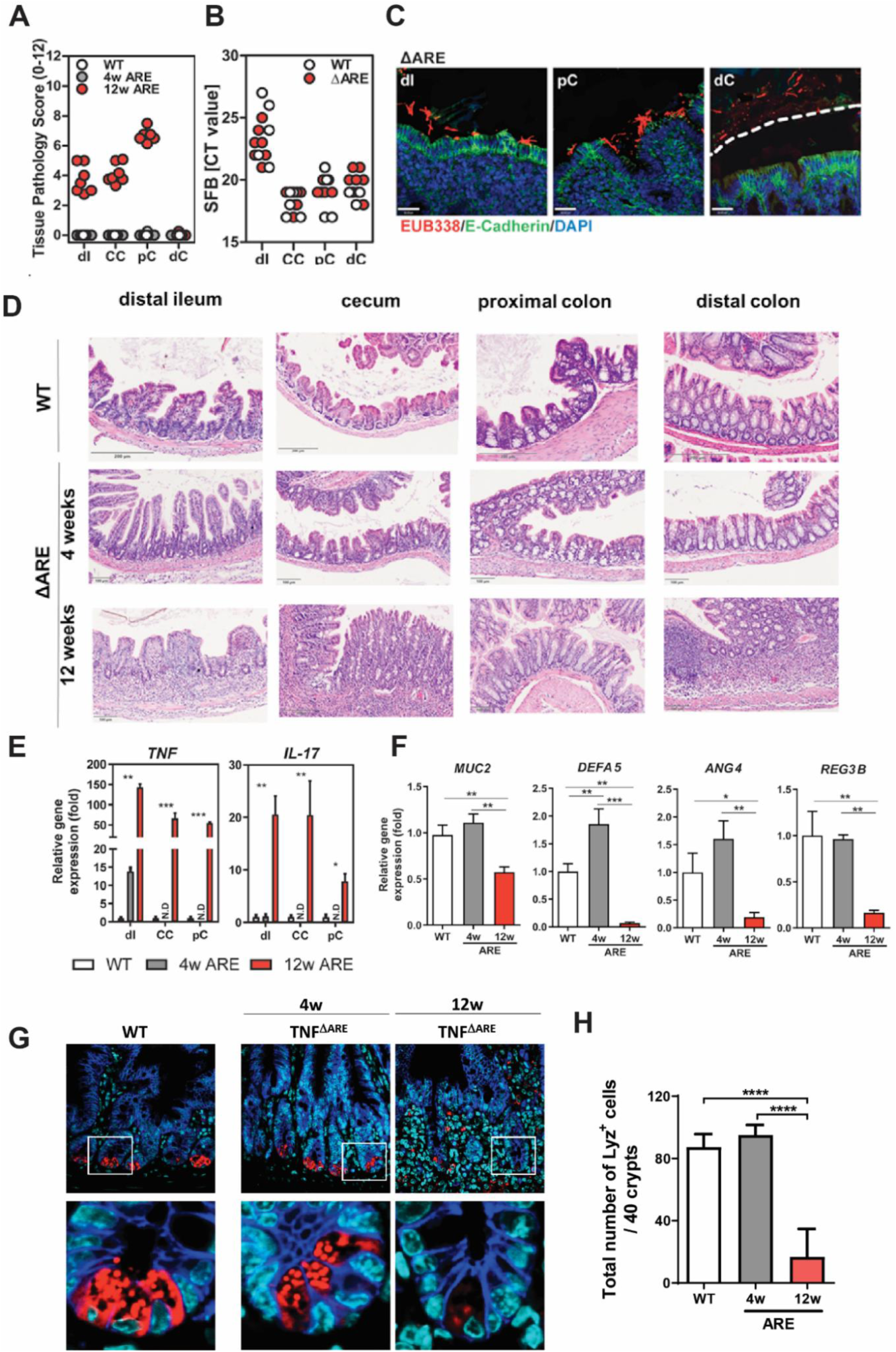
SFB mono-colonization causes CD-like inflammation in *Tnf* ^ΔARE^ mice. Germ-free *Tnf* ^ΔARE^ mice and littermate WT controls were monocolonized with SFB from birth. **(A)** Severity of inflammation was assessed by evaluating the tissue pathology score in tissue sections of distal ileum, cecum, and proximal and distal colon from 12 weeks-old WT mice as well as 4 weeks-old and 12 weeks-old *Tnf* ^ΔARE^ mice. **(B)** Quantitative analysis of SFB abundance in distal ileum, cecum, and proximal and distal colon of 12 weeks-old WT and *Tnf* ^ΔARE^ mice. **(C)** FISH by using the eubacterial probe EUB338 was performed on tissue sections from distal ileum and colon of 12 weeks-old *Tnf* ^ΔARE^ mice followed by immunostaining with E-Cadherin. DAPI was used for nuclear visualization. Dotted line indicates thickness of inner mucus layer. **(D)** H&E-stained tissue sections of 12 weeks-old *Tnf* ^ΔARE^ mice revealed profound intestinal inflammation in the compartments of distal ileum, cecum, proximal colon, but not jejunum and distal colon when compared to 12 weeks-old WT controls, but also 4 weeks-old *Tnf* ^ΔARE^ mice. Scale bar represents 200 µm. **(E)** Quantitative analysis of *Tnf a*nd *Il-17* transcript levels in mucosal tissue of distal ileum, cecum, and proximal colon from 4 and 12 weeks-old *Tnf* ^ΔARE^ mice and 12 weeks-old WT controls. Statistical significance was assessed by One-way analysis of variance (ANOVA) followed by Tukey test. *p<0.05; **p<0.01; ***p<0.001 was considered statistically significant. **(F)** Quantitative analysis of *Muc2*, *Defa5*, *Ang4*, and *Reg3b* transcript levels in mucosal tissue of distal ileum from 4 and 12 weeks-old *Tnf* ^ΔARE^ mice and 12 weeks-old WT controls. Statistical significance was assessed by One-way analysis of variance (ANOVA) followed by Tukey test. *p<0.05; **p<0.01; ***p<0.001 was considered statistically significant. **(G)** Immunofluorescence (IF) co-staining of Lysozyme (red) and E-cadherin (IEC borders, blue) counterstained with Dapi (nuclei, cyan) in ileal tissue sections from 4 weeks and 12 weeks mono-colonized WT and *Tnf* ^ΔARE^ mice (600x) lower panel: higher magnification of the indicated sections (3600x). **(H)** Graph represents quantification of the total number of Lysozyme positive Paneth cells per crypt based on IF staining. Statistical analyses were performed by One-way analysis of variance (ANOVA) followed by Tukey test.

### SFB fail to induce ileo-colonic inflammation in a colitis mouse model

We clearly demonstrated that SFB induces ileo-colonic inflammation in mono-colonized *Tnf* ^ΔARE^ mice. As SFB are known to attach to intestinal epithelial cells and potently stimulate host immune response, we next wanted to test host specificity to the mucosal-adherent SFB. For this purpose, we assessed the impact of SFB colonization on colitis development in IL10-deficient *(Il10^−/−^*129Sv) mice, known to develop Th1/Th17-related and microbiota-dependent colonic inflammation. In addition, we mono-colonized age matched *Tnf* ^ΔARE^ mice and WT controls for the same time frame. All groups were colonized at 8 weeks of age and analyzed 4 weeks later **(Figure 3A).** Intriguingly, SFB colonization did not induce inflammation in any of the intestinal compartments of *Il10^-/-^* mice. In contrast, *Tnf* ^ΔARE^ mice showed exacerbated inflammation in the proximal colon and in cecum tissue, and expectedly in the distal ileum tissue **(Figure 3B)**, as shown by microscopic analysis of H&E-stained tissue sections **(Figure 3D).** The presence of SFB in *Il10^-/-^* as well as *Tnf* ^ΔARE^ mice and their matching WT controls was confirmed using SFB-specific qPCR assay. Comparable SFB loads were observed in both mouse models **(Figure 3C).** To specifically examine changes in colon tissue morphology in both mouse models, we performed PAS/AB goblet cell staining in proximal colon tissue. The results showed significantly reduced numbers of PAS/AB positive mucin filled goblet cells in inflamed *Tnf* ^ΔARE^ mice, while *Il10^-/-^* mice remained unaffected **(Figure 3E and F).**

**Figure 3.**
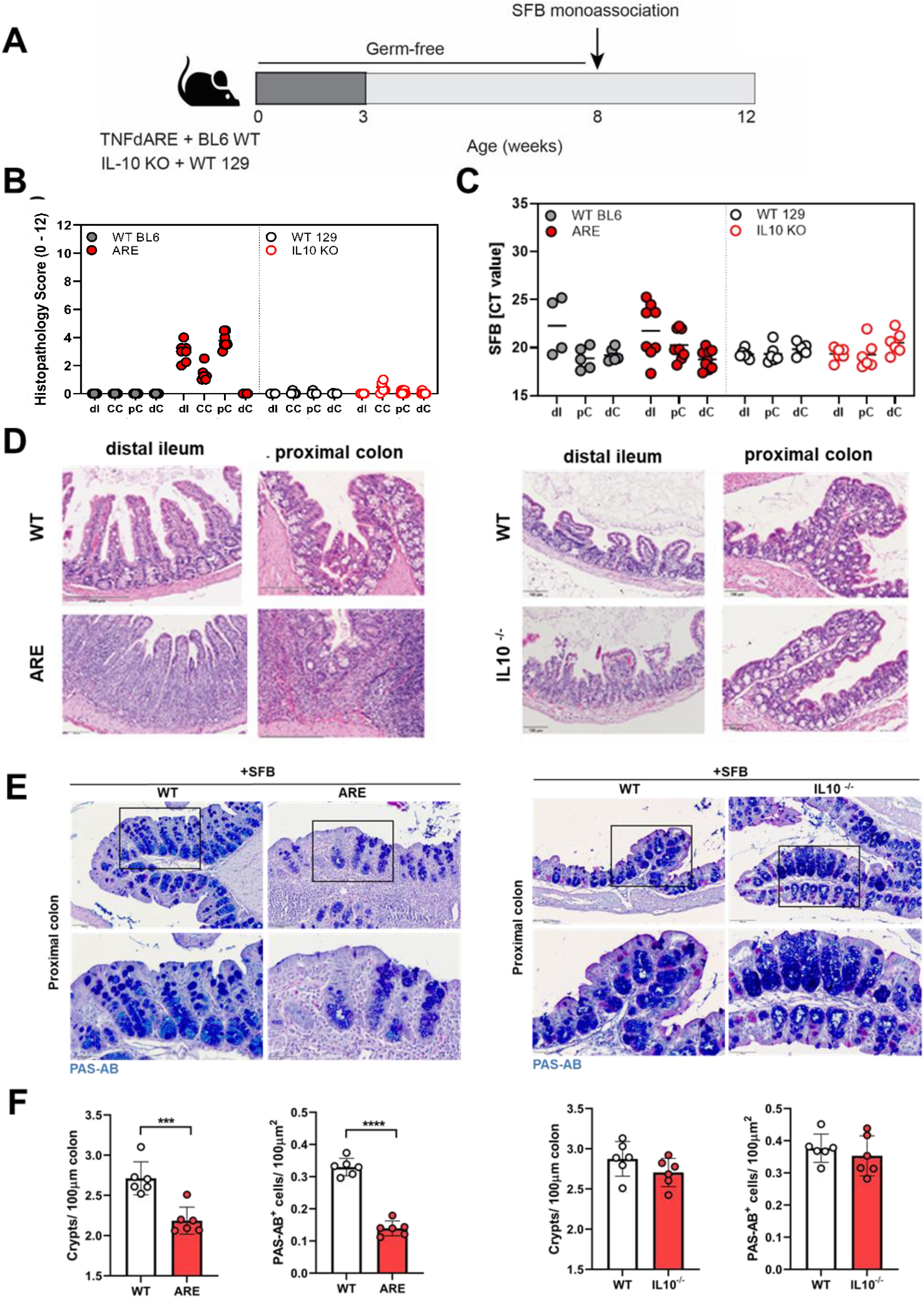
SFB fail to induce ileo-colonic inflammation in a colitis mouse model. **(A)** GF *Tnf* ^ΔARE,^ *Il10^-/-^* mice and WT controls were mono-colonized with SFB from 8 to 12 weeks of age. **(B)** Severity of inflammation was assessed by evaluating the tissue pathology score in tissue sections of cecum, proximal and distal colon from *Tnf* ^ΔARE^ mice and *Il10^-/-^* mice, respectively. **(C)** Quantitative analysis of SFB abundance in, cecum, proximal and distal colon of 12 weeks-old *Tnf* ^ΔARE^ mice and *Il10^-/-^* mice, respectively **(D)** H&E-stained tissue sections of *Tnf* ^ΔARE^ mice revealed intestinal inflammation in the compartments of distal ileum, cecum, proximal colon, compared to *Il10^-/-^* mice. Scale bar represents 200 µm. **(E, F)** PAS-AB goblet cell staining in proximal colon tissue sections from mono-colonized *Tnf* ^ΔARE^ and *Il10^-/-^*mice (400x). Lower panel represents higher magnification (1200x) of the indicated sections. Graph represents the quantification of goblet cells per crypt. Statistical analyses were performed by unpaired Student’s t-test.

### SFB pre-colonization is required to reactivate the pathogenicity of human-derived complex microbiota

We recently reported that the transplantation of fecal microbiota from inflamed CD patients successfully recreated the disease phenotype in recipient *Il10^-/-^* GF mice, while WT mice remained disease-free. Changes in immune cell profiles mirrored the level of tissue pathology and confirmed the transmissibility of disease activity in gnotobiotic mice (Metwaly et al., 2020). Moving forward, we wanted to test the capability of SFB to drive a patient-derived complex microbial community towards inflammation. To this end, we transplanted CD patient microbiota alone and in co-colonization with SFB into GF *Tnf* ^ΔARE^ mice. We colonized two groups of *Tnf* ^ΔARE^ mice and WT controls, either with fecal microbiota from CD patient only or in combination with SFB. Both groups were colonized at 8 weeks of age and assessed for tissue pathology at 12 weeks of age **(Figure 4A)**. Since SFB pathogenicity is seemingly dependent on the microbial community context besides the genetic susceptibility of the host, we wanted to test the capability of SFB to reactivate the pathogenic potential of a complex microbial community derived from CD patient. Consistent with our previous finding in *Il10^-/-^* (Metwaly et al., 2020), CD human remission microbiota transfer into *Tnf* ^ΔARE^ mice did not show any signs of inflammation or changes in ileal histopathology 4 weeks after colonization. However, SFB co-colonization induced inflammation in humanized *Tnf* ^ΔARE^ mice in contrast to mice colonized only with CD human microbiota **(Figure 4B, D).** SFB quantification using qPCR assay confirmed the colonization of SFB in the co-colonized mice (Ct values= 25.1±2.4), while they were undetectable (Ct values ≥ 35) in mice colonized only with human CD microbiota **(Figure 4C).** Consistently, SFB co-colonization resulted in reduced numbers of lysozyme positive Paneth cells **(Figure 4E, F).** To characterize microbial community shifts driven by the co-colonization with SFB, microbiota profiling of gut content showed a clear separation between *Tnf* ^ΔARE^ mice colonized with CD human microbiota alone or in combination with SFB. These changes were characterized by an overabundance of *Alistipes* and *Bilophila* (**Figure 4G, H)**. Interestingly and like the microbial shifts observed in humanized SFB co-colonized mice, *Alistipes* showed to be one of the most significantly differential taxa in *Tnf* ^ΔARE^ mice under SPF conditions. We next sought to examine the presence of SFB in human mucosal biopsies and screened 412 ileal and colonic biopsies derived from 3 independent cohorts of IBD patients (**Supplementary Figure 3A)**. We used a PCR approach and the primers previously published by Jonsson (Jonsson, 2013) and by Snel (Snel et al., 1995) and the newly designed primer set (unpublished data, Pamela Schnupf). In addition, we used SFB cloned DNA and 2 SFB human biopsies positive for SFB as positive control (**Supplementary Figure 3B)**. In these experiments, 0.3 or 3 ng of input DNA was used, and gene amplification of the newly designed primer set was confirmed by the detection of the expected 200bp PCR product via gel electrophoresis (**Supplementary Figure 3C)**. While only 27/412 samples showed CT values below than 35, PCR yielded random amplification with SFB identity not verified. The second and newly designed primer set showed specific amplification and the identity of amplified PCR products with SFB was verified by DNA sequencing and alignment to the positive control (**Supplementary Figure 3D-G)**. Nevertheless, none of the tissue biopsies that proved to be positive using the first primer-set showed to be positive for SFB.

### EEN-like purified diet eradicates SFB and prevents CD-like ileo-colonic inflammation in *Tnf* ^ΔARE^ mice

To simulate the protective effect of exclusive enteral nutrition (EEN) therapy in humans, we fed *Tnf* ^ΔARE^ mice and WT matching control EEN-like purified diet (PD) directly after weaning and characterized bacterial and host responses after 10 weeks of feeding. Our data showed that PD completely inhibited the development of intestinal inflammation in *Tnf* ^ΔARE^ mice and correlated with inflammation severity **(Suppl. Figure 4A)** . The diet-mediated protection was also associated with substantial changes in community structure **(Suppl. Figure 4B,C)**.

To investigate the direct influence of diet on SFB colonization and on intestinal inflammation, we fed SFB mono-colonized *Tnf* ^ΔARE^ mice with chow or EEN-like purifiet diet **(Figure 5A)**. In the first experiment, WT and *Tnf* ^ΔARE^ mice were colonized with SFB at 8 weeks of age and were fed with chow diet until the end of the experiment. In the second and third experiments, the mice were colonized with SFB at 8 weeks of age, and were exposed to purified PD one week before or after SFB colonization (7 weeks or 9 weeks of age, respectively). Expectedly, SFB quantification using qPCR assay confirmed the presence of SFB in ileal, cecal and colonic content of both WT and *Tnf* ^ΔARE^ SFB mono-colonized mice fed with chow diet (Ct values= 21.5±2.5). On the other hand, SFB were undetectable (Ct values ≥ 35) in SFB-mono-colonized mice fed with PD before or after SFB colonization **(Figure 5C).** Histopathological evaluation revealed inflammation in the distal ileum and proximal colon of *Tnf* ^ΔARE^ mice fed with chow diet, while mice fed with PD showed no signs of inflammation **(Figure 5B, Suppl. Figure 4D).** Gene expression analysis on tissue sections from distal ieum showed significanly elevated expression of *Tnf* and *Il-17* in *Tnf* ^ΔARE^ mice mono-colonized with SFB and fed with chow diet compared to those fed with PD (**Suppl. Figure 4E).** Consistently, a significantly reduced number of lysozyme positive Paneth cells was observed in *Tnf* ^ΔARE^ mice on chow diet compared to mice on PD **(Figure 5D).** To visualize SFB in tissue sections of *Tnf* ^ΔARE^ mice under different dietary conditions, we performed fluorescence in situ hybridization (FISH) by staining proximal colon sections with bacteria domain-specific (EUB338) probes. Microscopic examination showed growth of filamentous bacteria close to the intestinal epithelium in *Tnf* ^ΔARE^ mice on chow diet, while they were not visible in *Tnf* ^ΔARE^ mice under PD (**Figure 5E).**

**Figure 5.**
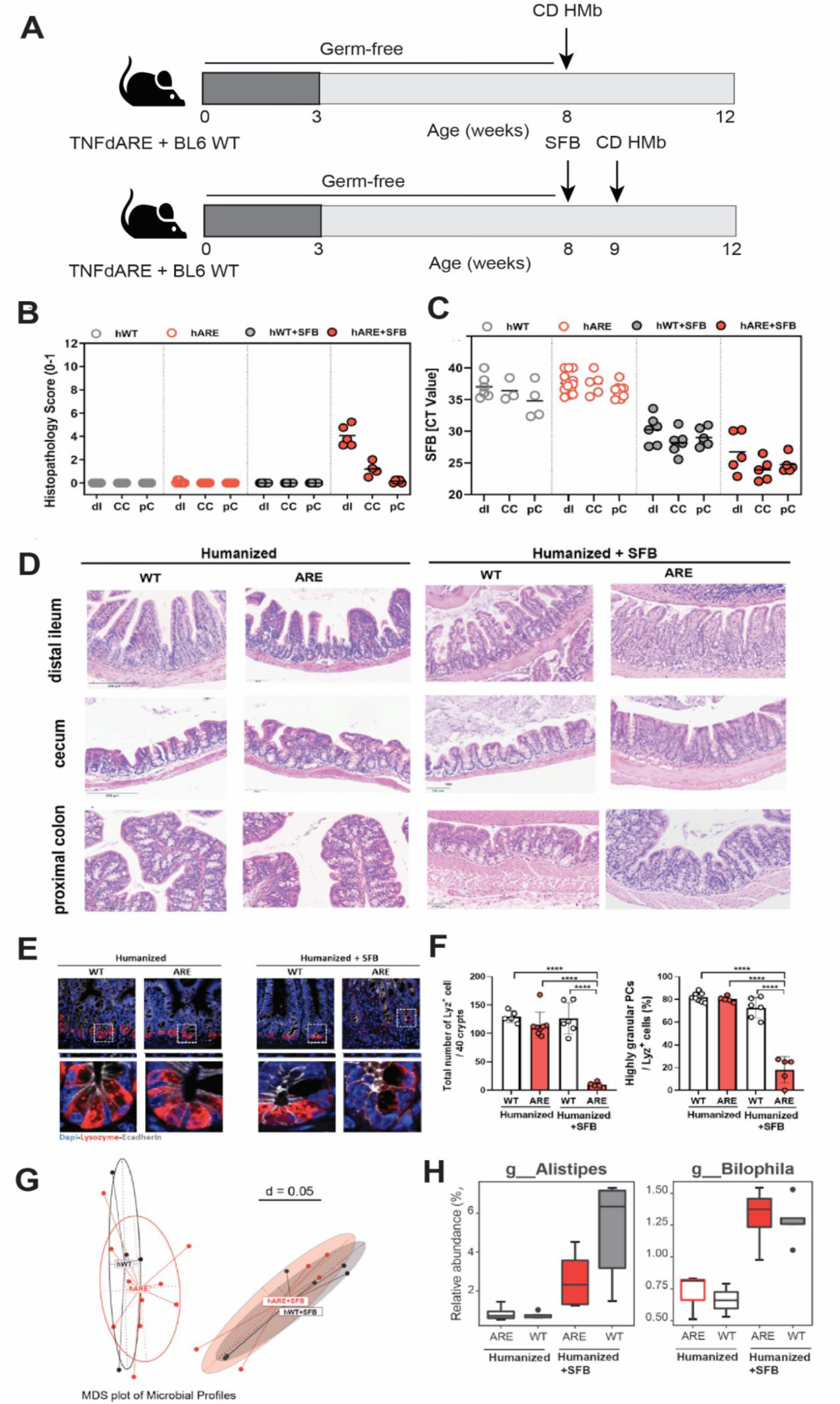
SFB initiate inflammation in humanized *Tnf* ^ΔARE^ mice. **(A)** *Tnf* ^ΔARE^ mice and WT controls were colonized either with fecal microbiota from CD patient only or in combination with SFB. Both groups were colonized at 8 weeks of age and assessed for tissue pathology at 12 weeks of age**. (B)** Severity of inflammation was assessed by evaluating the tissue pathology score in tissue sections of distal ileum, cecum and proximal colon of humanized or SFB + humanized *Tnf* ^ΔARE^ mice. **(C)** Quantitative analysis of SFB abundance in humanized or SFB + humanized *Tnf* ^ΔARE^ mice. **(D)** H&E-stained tissue sections of *Tnf* ^ΔARE^ mice revealed enhanced intestinal inflammation in the distal ileum and to a lower degree in cecum tissue of humanized *Tnf* ^ΔARE^ mice co-colonized with SFB. **(E, F)** Immunofluorescence (IF) co-staining of Lysozyme (red) and E-cadherin (IEC borders, blue) counterstained with Dapi (nuclei, cyan) in ileal tissue sections from *Tnf* ^ΔARE^ and WT mice colonized with human CD microbiota alone (hARE, hWT) or in co-colonization with SFB (hARE+SFB, hWT+SFB) (600x) lower panel: higher magnification of the indicated sections (3600x). Graph represents quantification of the total number of Lysozyme positive Paneth cells per crypt based on IF staining. Statistical analyses were performed by One-way analysis of variance (ANOVA) followed by Tukey test. **(G)** Beta-diversity analysis of bacterial community profiles in *Tnf* ^ΔARE^ and WT mice colonized with human CD microbiota alone (hARE, hWT) or in co-colonization with SFB (hARE+SFB, hWT+SFB) **(H)** Significantly increased percentage relative abundance of *Alistipes* and *Blautia* in humanized *Tnf* ^ΔARE^ and WT mice co-colonized with SFB (hARE+SFB, hWT+SFB).

**Figure 6.**
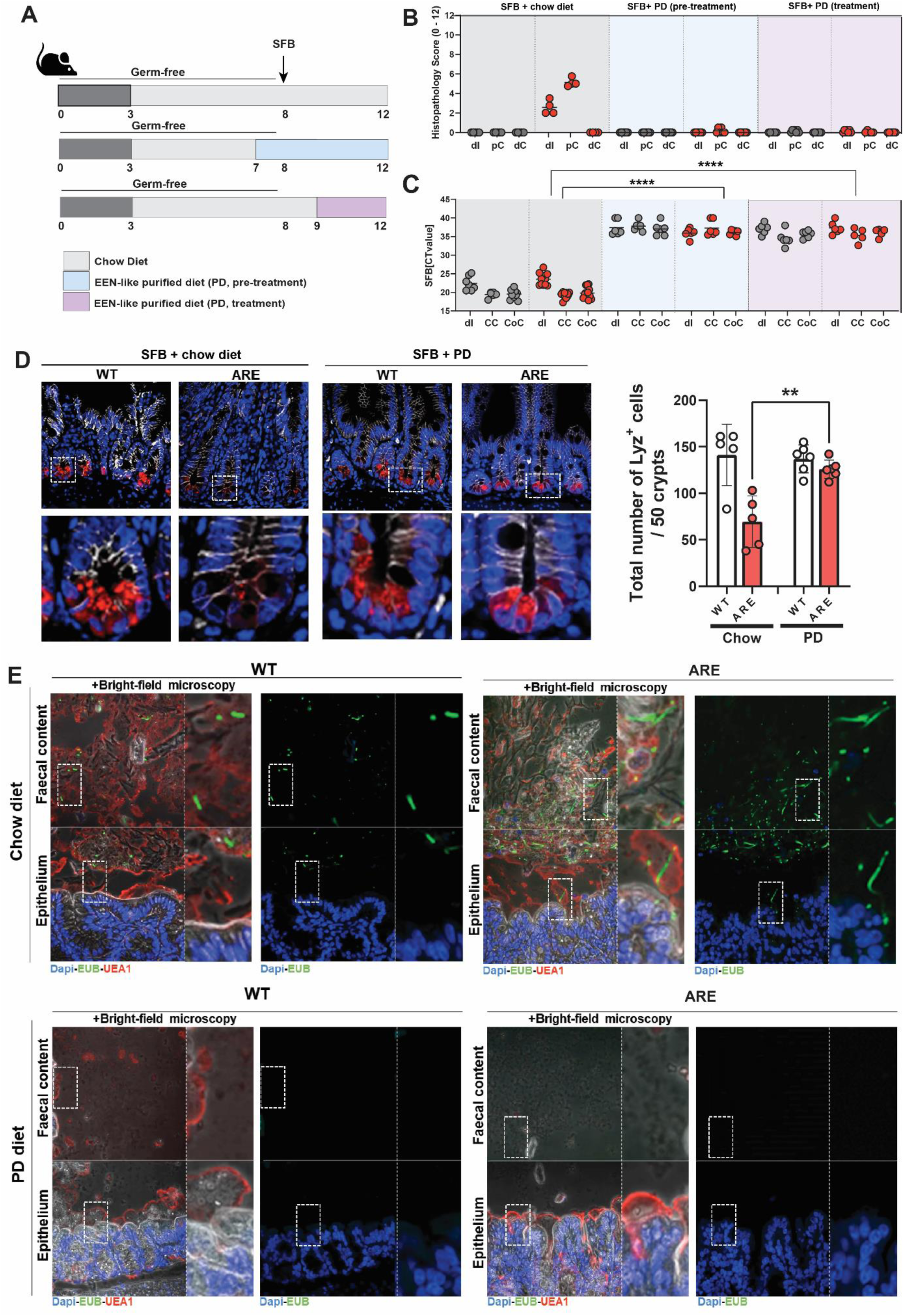
EEN-like PD eradicates SFB and prevents CD-like ileo-colonic inflammation in *Tnf* ^ΔARE^ mice. **(A)** Germ-free *Tnf* ^ΔARE^ mice and littermate WT controls were mono-colonized with SFB at 8 weeks of age and exposed to chow or EEN-like PD **(B)**. Severity of inflammation was assessed by evaluating the tissue pathology score in tissue sections of distal ileum, proximal and distal colon in WT and *Tnf* ^ΔARE^ mice. **(C)** Quantitative analysis of SFB abundance in ileal, cecal, and colonic contents of chow-fed or PD-fed WT and *Tnf* ^ΔARE^ mice **(D)** Immunofluorescence (IF) co-staining and quantification of Lysozyme (red) and E-cadherin (IEC borders, blue) counterstained with Dapi (nuclei, cyan) in ileal tissue sections from mono-colonized and chow or PD-fed WT and *Tnf* ^ΔARE^ mice (600x) lower panel: higher magnification of the indicated sections (3600x). **(E)** FISH by using the eubacterial probe EUB338 was performed on tissue sections from distal ileum and colon of chow-fed or PD-fed WT and *Tnf* ^ΔARE^ mice followed by immunostaining with E-Cadherin. DAPI was used for nuclear visualization. Dotted line indicates thickness of inner mucus layer.

## DISCUSSION

Accumulating evidence suggests a crucial role of the intestinal microbiota in the onset and progression of inflammatory bowel diseases (Morgan et al., 2012; Nagao-Kitamoto and Kamada, 2017; Sartor and Wu, 2017). Previous reports identified a few pathogenic bacteria that enrich in dysbiotic microbial communities and lead to disease development (Darfeuille-Michaud et al., 2004; Prindiville et al., 2000). In the present study, we aimed at identifying the causal microbial cues responsible for inducing or modulating Crohn’s disease-like inflammation in *Tnf* ^ΔARE^ mice. Intriguingly, histological analysis of ileal tissue sections clearly revealed the detection of filamentous bacteria in inflamed responder mice, closely resembling the commensal SFB. SFB are well known to induce the maturation of Th17 cell-derived immune responses, which are highly implicated in the pathogenesis of IBD (Gálvez, 2014). However, assessment of disease development kinetics in *Tnf* ^ΔARE^ mice showed that both *Tnf* ^ΔARE^ mice as well as littermate WT controls harbour significantly higher SFB loads at the age of 3-4 weeks, which declined below detectable levels in adult mice, under non-inflammatory conditions. Similar observations were reported, describing bacterial alterations as an ongoing physiological process in line with the intestinal mucus layer maturation upon bacterial colonization (Johansson et al., 2015). Interestingly, we showed that in adult *Tnf* ^ΔARE^ mice, the abundance of SFB strongly correlated with the severity of ileal inflammation, suggesting that SFB growth is driven by changes in the bacterial community composition, subsequently leading to inflammation.

To investigate whether SFB can persist in a maturate immune system, and whether they are capable of driving changes in a complex intestinal bacterial community towards inflammation. Interestingly, the introduction of SFB in SPF-housed 8 weeks old *Tnf* ^ΔARE^ mice significantly enhanced the abundance of *Alistipes* species that has been previously reported to trigger colitis and tumour growth in *Il-10*-deficient mice (Moschen et al., 2016). Under physiological conditions, the growth of *Alistipes* is suggested to be regulated by iron availability (Goetz et al., 2002). In this context, work from our group showed that the induction of inflammation in *Tnf* ^ΔARE^ mice seemed to be highly dependent on the availability of dietary iron since mutant mice fed with low-iron diet remained healthy and showed no signs of inflammation (Werner et al., 2011). Conclusively, we suggested that *Alistipes* might be involved in the onset of CD-like ileitis in *Tnf* ^ΔARE^ mice, but mono-colonization of mice with *Alistipes* was not sufficient to trigger inflammation. Since SFB is also known to utilize iron as an energy source for growth, we thus assumed that propagating SFB are rather improving than impairing the growth conditions for *Alistipes* by possibly enhancing substrate availability (Sczesnak et al., 2011). We next proposed that most likely inflammation in *Tnf* ^ΔARE^ mice is triggered by the mere presence of SFB. Here we showed that the mono-colonization of *Tnf* ^ΔARE^ mice with SFB from birth and for 12 weeks could induce severe inflammation which expanded to the colon, thereby demonstrating for the first time the pathogenic role of these commensal bacteria in the intestine. In general, *Tnf* is upregulated in *Tnf* ^ΔARE^ mice upon microbial colonization. Considering our findings, enhanced *Tnf* transcript levels are not sufficient to induce inflammation in *Tnf* ^ΔARE^ mice but cause changes in the immune responses triggered by SFB. While SFB are well known to impact the differentiation of gut mucosal Th17 cells, SFB-mono-colonized *Tnf* ^ΔARE^ mice additionally showed significantly expanded populations of IFN-expressing CD4+ cells and neutrophilic granulocytes as well as an impaired epithelial antimicrobial defence of the intestine. In this context, Flannigan et al. (2017) described the recruitment of neutrophils into the ileum upon SFB-mediated IL-17A expression as a control mechanism limiting SFB expansion in the gut (Flannigan et al., 2017).

Consistent with these findings, high SFB loads have been reported in a subset of UC patients, suggesting a potential role for SFB in driving intestinal inflammation in human (Finotti et al., 2017). Nevertheless, it needs to be acknowledged that contradictory data exist regarding the presence of SFB in humans. An early report proved the presence of a filamentous organism, potentially being SFB, on the ileal mucosa of a human subject using light microscopy (Klaasen et al., 1993). In 2011, a large-scale study by Scyesnak *et al*. concluded that SFB are not present in humans based on metagenomic datasets search (Sczesnak et al., 2011). These findings were contradicted by a report showing the presence of SFB in human feces from many individuals. Arguably, the 16S rRNA gene sequences corresponding to human SFB in this work showed to cluster together with mouse SFB sequences (Yin et al., 2013). In a recent work, Jonsson *et al*. published the draft genome sequence of human-adapted representative of SFB in a human ileostomy sample (Jonsson et al., 2020) which was validated by screening metagenomic datasets and identifying individuals carrying sequences identical to the new SFB genome. In the present study, we investigated the presence of SFB in 412 tissue biopsies collected from three independent cohorts of adult and paediatric IBD patients. We screened and analysed the mucosal biopsies using previously published (Finotti et al., 2017) and newly designed human SFB specific primers, based on the recently published SFB human genome sequence (Jonsson et al., 2020). Interestingly, SFB were absent in ileal and colonic mucosal biopsies from IBD patients with active or inactive disease To test the capability of human gut microbiota to drive inflammation in *Tnf* ^ΔARE^ mice as we and others showed previously using *Il-10* deficient mice (Metwaly et al., 2020; Nagao-Kitamoto et al., 2016), we colonized *Tnf* ^ΔARE^ mice with faecal microbiota derived from an IBD patient. Surprisingly and in contrary to *Il-10* deficient mice, the same patient microbial communities failed to induce inflammation in *Tnf* ^ΔARE^ mice. Considering the strong immunomodulatory functions of SFB, they can pose a synergistic effect with other commensal bacteria to drive beneficial or detrimental effects on the host physiology (Ericsson et al., 2015). Investigating the capability of SFB to reactivate the pathogenic potential of human-derived microbiota through dual colonization of *Tnf* ^ΔARE^ with SFB and IBD patient-derived gut microbiota led to ileal inflammation together with reduced numbers of functional Paneth cells in *Tnf* ^ΔARE^ mice, suggesting that SFB contribute to microbial community and regulatory immune response modulation, leading to inflammation. Notably, the beneficial modulatory effect of SFB in a complex microbial community was shown in a recent study, where co-colonization with SFB abolished the murine norovirus (MNV)-induced colitis in Altered Schaedler Flora (ASF) colonized mice. In this context, SFB colonization enhanced the expression of pro-inflammatory cytokines and antimicrobial peptides, leading to inhibition of inflammation (Bolsega et al., 2019).

Intriguingly, prior evidence showed that dietary composition is an important determinant of the presence of SFB in the mouse small intestine. In an early study, the exclusive feeding in mice with milk powder showed to inhibit the growth and colonization of SFB when compared to complete laboratory mouse diets feeding (Klaasen et al., 1991). In this study, the authors used twenty-one different purified diets as mouse feed to identify the macronutrients that might be responsible for SFB growth and colonization. Interestingly, none of the purified diets allowed SFB growth, however the critical dietary cues responsible for these observations were not identified (Klaasen et al., 1991). These findings suggest an important role of diet in modulating SFB growth and hence SFB-mediated inflammation.

The best and most prominent evidence for the ability of specific nutritional intervention in IBD treatment stems from the use of exclusive enteral nutrition (EEN) as induction therapy for pediatric CD. Changes in microbiota composition upon EEN therapy have been recently demonstrated (Schwerd et al., 2016), but the causality behind the success of this dietary intervention is not understood. EEN is a therapeutic nutrition confined in most cases to liquid fiber-free enteral formulas of typically synthetic or semi-synthetic composition. Here, we showed that consistent with the impact of EEN on disease activity in pediatric CD patients, the use of a purified experimental diet that is low in fiber content completely inhibited the development of CD-like ileitis in *Tnf* ^ΔARE^ mice under SPF housing conditions. These effects were associated with substantial changes in community structure. To elucidate the direct effect of the purified diet on SFB growth and mediated inflammation, we performed mono-colonization studies and showed that the pre-treatment or subsequent treatment of SFB mono-colonized *Tnf* ^ΔARE^ mice with the EEN-like purified diet completely prevented or abolished SFB, respectively. One of the main differences between PD or EEN and chow diet is the presence of dietary fibers. Interestingly, in a previously published dietary intervention study, SFB were detected in two ileostomy samples from one patient during high-fiber diet intake, while they were not present in samples from the same patient during low-fiber diet intake. Based on the observations that SFB are present at higher abundances during weaning in many animals which could be because of the introduction of solid food or dietary fiber instead of mother’s milk, we hypothesized that fiber could be the critical determinant of SFB propagation in *Tnf* ^ΔARE^ mice. Parallel to these findings, an early study in 1992 demonstrated that fibre-rich beans lead to an increased SFB colonisation in mice, when added to a natural diet (Klaasen et al., 1992). Taken together, these data suggest the importance of dietary fibre availability for SFB growth in a mouse model for CD-like ileitis and proposes a plausible mechanism explaining the effectiveness of exclusive enteral nutrition in treating patients with IBD.

In conclusion, this study aimed to identify the causal microbial cues responsible for CD-like inflammation and to dissect the protective role of diet in gnotobiotic mouse models providing a causal link between pathobiont expansion and CD-like ileo-colonic inflammation. We identified a novel pathogenic role of SFB in driving severe ileo-colonic inflammation in murine models, while SFB was absent in mucosal biopsies from IBD patients. Simulating the protective effect of EEN by feeding SFB mono-associated *Tnf* ^ΔARE^ mice chow diet or EEN-like purified diet showed that the pre-treatment or the subsequent treatment of SFB mono-colonized *Tnf* ^ΔARE^ mice with EEN-like purified diet prevented SFB colonization and completely abolished SFB-mediated inflammation, suggesting that fiber is a critical determinant of pathobiont expansion and ileal inflammation in this model. Collectively, these findings provide clear evidence for the importance of dietary fiber availability for SFB growth in a mouse model for CD-like ileitis and propose a plausible mechanism explaining the mechanism of action EEN efficacy in children with IBD.

## Funding

Funded by the Deutsche Forschungsgemeinschaft (DFG, German Research Foundation) – project number 395357507 (SFB 1371, Microbiome Signatures).

## Competing interests

None

The authors declare no competing interests.

---

**Supplemental Figure 1.**
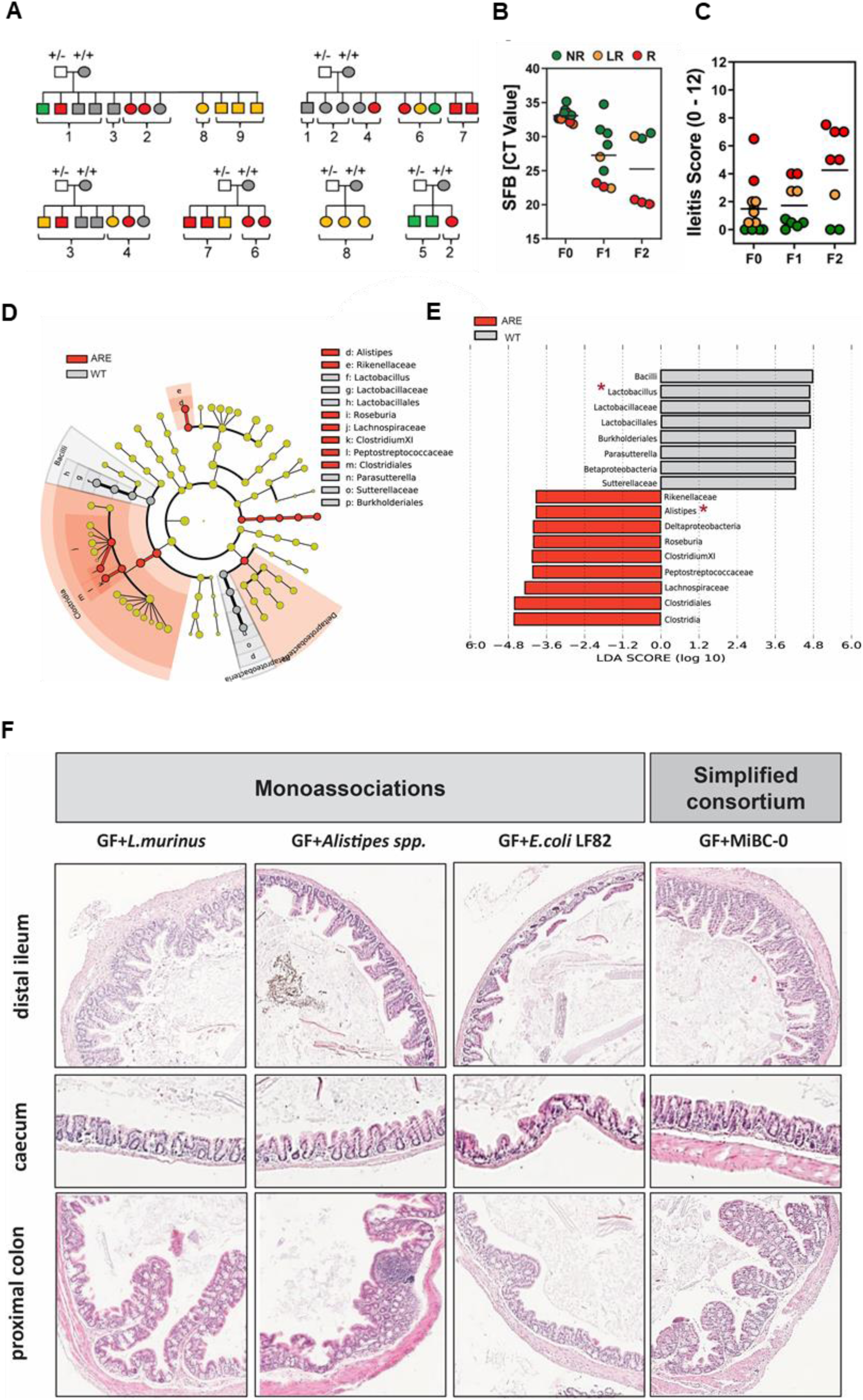
SFB abundance correlates with ileitis severity in SPF-house *Tnf* ^ΔARE^. **(A)** Litter and cage effect on ileitis development in SPF-housed *Tnf* ^ΔARE^ mice. Squares represent males; circles indicate females. Green, orange, and red symbols indicate *Tnf* ^ΔARE^ mice at 18-week endpoint with no (score 0), low (score <4) and high (score >4) ileitis histopathological score, respectively; grey symbols indicate WT littermates that do not develop ileitis; white symbols indicate male mice of unknown ileitis status. Each cage is delineated with the brackets below and a cage number. **(B)** Quantitative Analysis of SFB in recolonized *Tnf* ^ΔARE^ mice (F0, F1, F2). Color-code represents the severity of inflammation as described above. CT value >30 is regarded as non-specificity threshold. **(C)** Ileitis scores of recolonized 18-weeks-old *Tnf* ^ΔARE^ mice including 2 breeding generations (F0, F1, F2). Mice are color-coded based on inflammation severity with green (score 0); orange (score <4) (orange); and red (score >4). **(D)** Cladogram obtained from Linear discriminant analysis effect size (LEfSe) analysis of taxonomic profiling using 16S rRNA gene sequencing of intestinal microbiota in WT and *Tnf*^ΔARE^ mice. **(E)** Comparison of relative abundance of bacterial genera between WT and *Tnf* ^ΔARE^ mice using LEfSe analysis. Taxa meeting an LDA significant threshold 2 are shown, taxa enriched in *Tnf* ^ΔARE^ mice (red) and taxa enriched in WT mice (blue). **(F)** Representative H&E-stained tissue sections from distal ileum, caecum, and proximal colon of *Tnf* ^ΔARE^ mice colonized with single bacterial strains (*Alistipes*, *Lactobacillus murinus* and *E. coli LF82*) or with MIBAC, a minimal consortium of 7 bacterial strains showing no signs of inflammation.

**Supplementary Figure 2.**
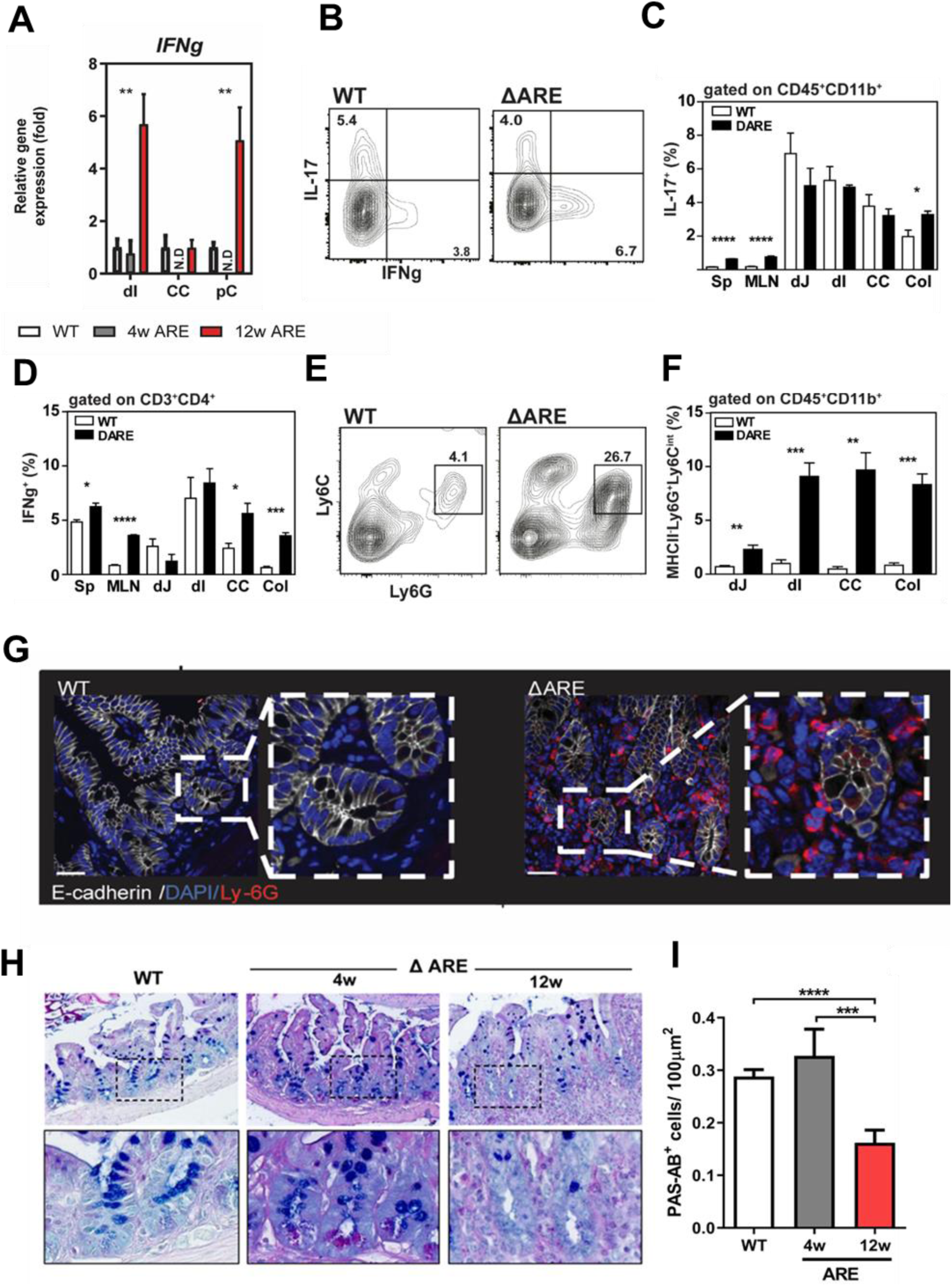
Enhanced numbers of IL-17- and IFNg-expressing CD4-positive cells as well as neutrophilic granulocytes in inflamed mucosal tissue of *Tnf* ^ΔARE^ mice. Immune cells were isolated from spleen, MLN, and intestinal mucosa of jejunum, ileum, cecum, and colon of 12 weeks old *Tnf* ^ΔARE^ mice (n = 3) and WT controls (n = 4). (A) Quantitative analysis of *Ifng* transcript levels in mucosal tissue of distal ileum, cecum, and proximal colon from 4 and 12 weeks-old *Tnf* ^ΔARE^ mice and 12 weeks-old WT controls. Statistical significance was assessed by xy. *p<0.05; **p<0.01; ***p<0.001 was considered statistically significant. (B) Gating strategy used for the assessment of IL-17- and IFNg-expressing CD3^+^CD4^+^ immune cells. Discrimination of live and dead cell population was performed by applying specific fluorescent dye. Percentage of IL-17- (C) and IFNg- (D) expressing cells was determined by setting gates within the CD3^+^CD4^+^ population. (E) Gating strategy used for the assessment of neutrophilic granulocytes. Discrimination of live and dead cell population was performed by applying specific fluorescent dye. Population of neutrophils was identified by applying antibodies against CD45, CD11b, MHCII, Ly6C, and Ly6G. (F) Percentage of neutrophilic granulocytes was assessed within the CD45^+^CD11b^+^ population by gating for MHCII^-^ Ly6C^int^Ly6G^high^ cells. (G) Immunofluorescence staining of tissue sections from distal ileum of WT and *Tnf* ^ΔARE^ mice applying Ly6G and E-Cadherin. DAPI was used for nuclear visualization. (H) PAS-AB goblet cell staining in ileal tissue sections from 4 weeks and 12 weeks mono-colonized WT and *Tnf* ^ΔARE^ mice (400x). Lower panel represents higher magnification (1200x) of the indicated sections. Graph represents the quantification of goblet cells (x1000) in 1µm2 selected crypt area. Statistical analyses were performed by One-way analysis of variance (ANOVA) followed by Tukey test. *p<0.05; **p<0.01; ***p<0.001 was considered statistically significant.

**Supplementary Figure 3.**
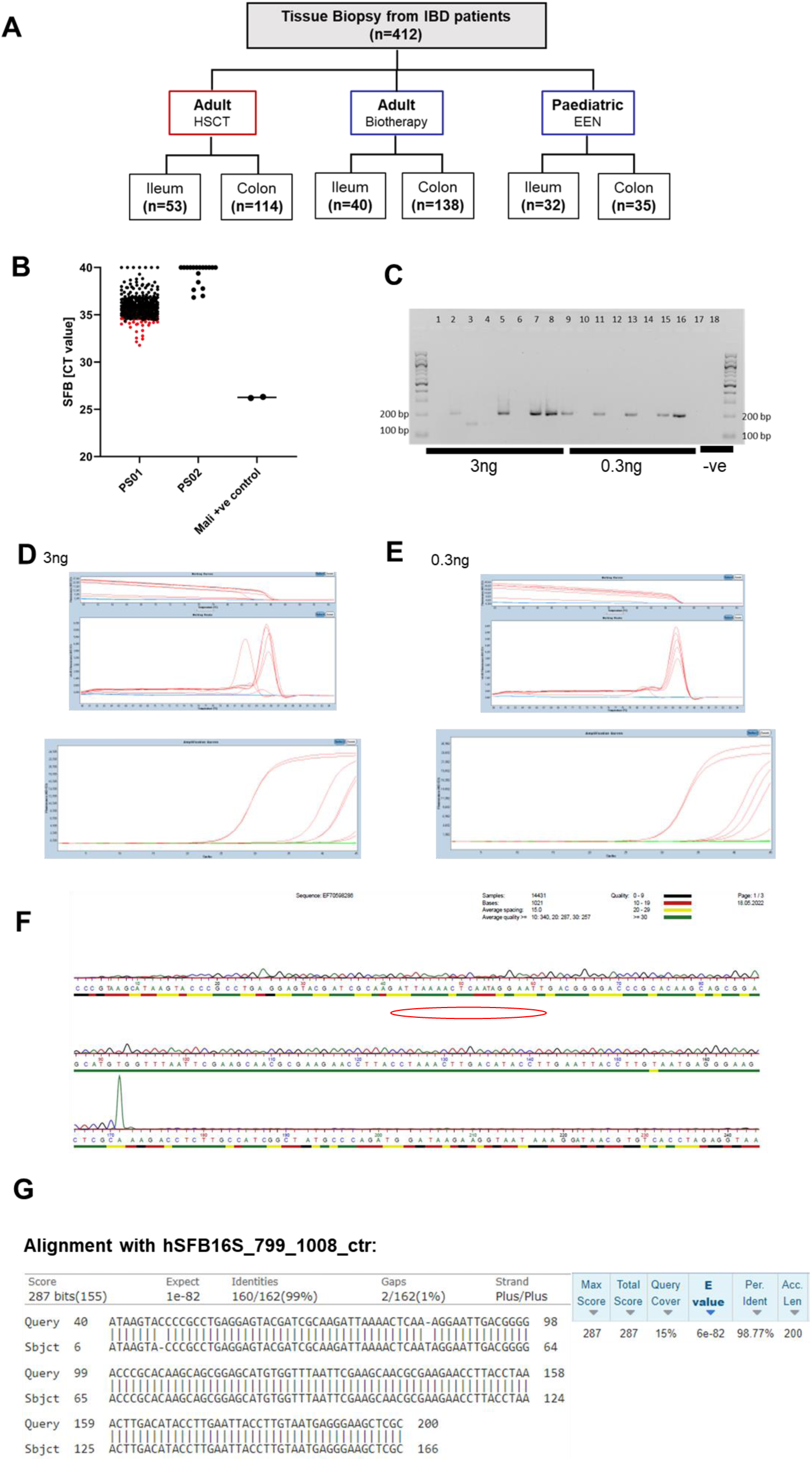
**(A)** Human cohorts screened for the presence of SFB **(B)** Quantitative analysis of SFB abundance in tissue biopsies using two primer sets (**C)** Expected 200 bP PCR products (**D, E)** Melting curves and CT values **(F, G)** Alignment to control human SFB sequence yielding 99% similarity

**Supplementary Figure 4.**
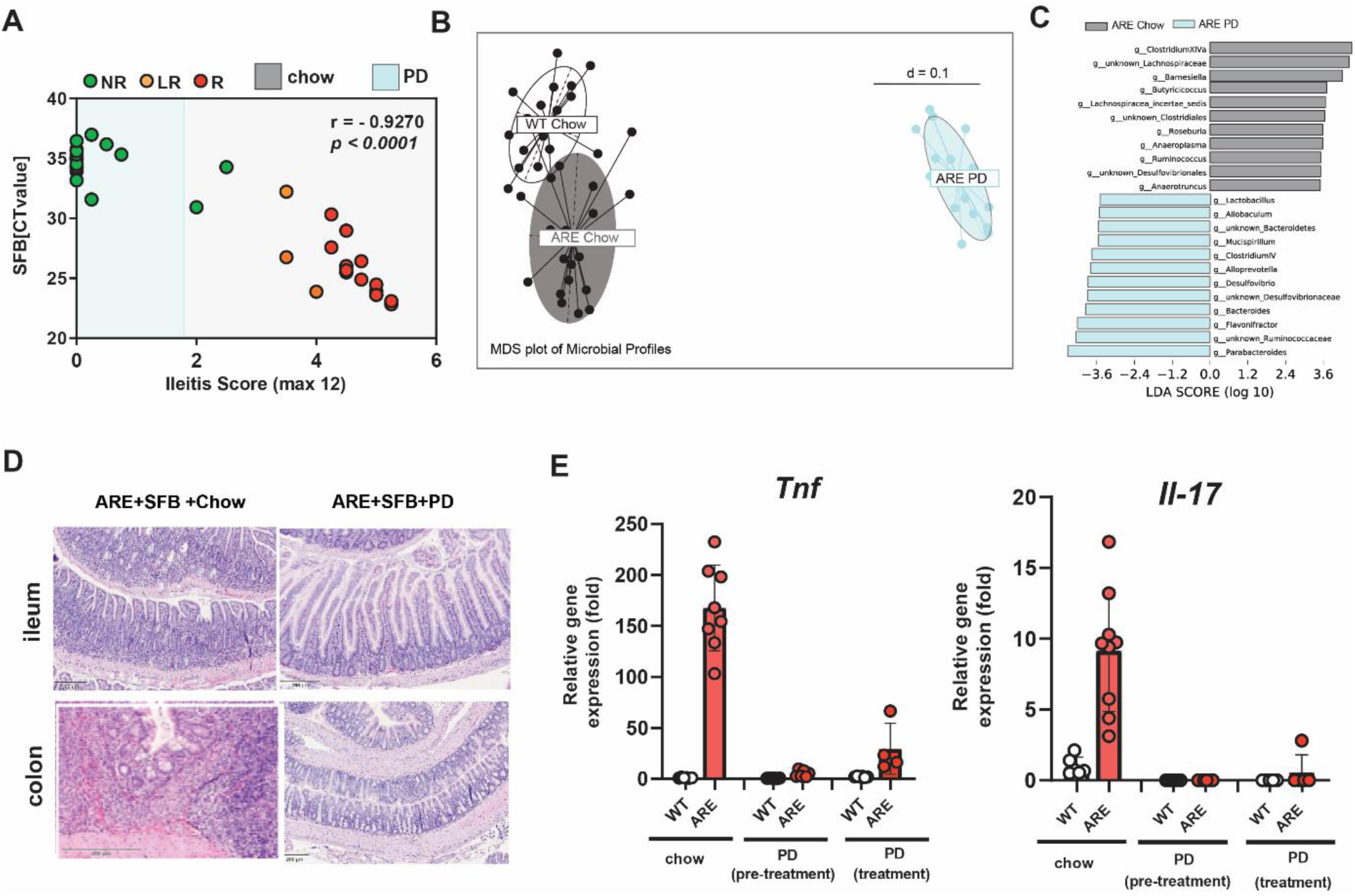
**(A)** Correlation analysis between Ileitis score and SFB abundance in chow-fed or PD-fed Specific pathogen-free (SPF) *Tnf* ^ΔARE^ mice and. **(B)** MDS analysis show separation of caecal bacterial communities according to diet among chow-fed WT, chow-fed or PD-fed *Tnf* ^ΔARE^ mice (**C)** Comparison of relative abundance of bacterial genera between *Tnf* ^ΔARE^ mice fed with chow or PD under SPF conditions, using LEfSe analysis. Taxa meeting an LDA significant threshold 2 are shown **(D)** Histopathology section of ileal and colonic tissue in ileal and colonic tissue of *Tnf* ^ΔARE^ mice mono-colonized with SFB and fed with chow or PD **(E)** Quantitative analysis of *Tnf a*nd *Il-17* transcript levels in mucosal tissue of distal ileum from *Tnf* ^ΔARE^ mice fed with chow or PD. PD: purified diet.

## Notes

### Competing Interest Statement

The authors have declared no competing interest.

