## Supplementary information for "Diet prevents the expansion of segmented filamentous bacteria and ileo-colonic inflammation in a model of Crohn’s disease"

**Supplementary Table S1:** Composition of chow, purified and fiber-rich diet

| | Chow Diet | PD | FRD |
|---|---|---|---|
| Carbohydrates [E%] | 61 | 64 | 61-64 |
| Fat [E%] | 12 | 13 | 12-13 |
| | soybean oil | soybean oil | soybean oil |
| Protein [E%] | 27 | 23 | 23-27 |
| Fiber [g/kg] | Not specified; prob. 200-300 | 50 | 200-300 |
| | Cellulose, hemicellulose, lignin (not specified) | Cellulose | Mixed Complex (simulating fiber content in Chow Diet) |
| Simple Sugar [g/kg] | Not specified and not added | 60 | 60 |

**Supplementary Table S2:** Description and metadata of patient mucosal biopsy samples screened for SFB presence.

| No | Sample-ID | Cohort | Phenotype | Location |
|---|---|---|---|---|
| **1** | B025 | HSCT | CI | ASCENDING COLON |
| **2** | B053 | HSCT | CI | ASCENDING COLON |
| **3** | B087 | HSCT | CIA | ASCENDING COLON |
| **4** | B148 | HSCT | CH | ASCENDING COLON |
| **5** | B066 | HSCT | CIA | DESCENDING COLON |
| **6** | B074 | HSCT | CII | DESCENDING COLON |
| **7** | B078 | HSCT | CH | DESCENDING COLON |
| **8** | B092 | HSCT | CH | DESCENDING COLON |
| **9** | B097 | HSCT | CIA | DESCENDING COLON |
| **10** | B024 | HSCT | IH | ILEUM |
| **11** | B044 | HSCT | IH | ILEUM |
| **12** | B052 | HSCT | IH | ILEUM |
| **13** | B067 | HSCT | IH | ILEUM |
| **14** | B091 | HSCT | III | ILEUM |
| **15** | B096 | HSCT | IH | ILEUM |
| **16** | B147 | HSCT | IH | ILEUM |
| **17** | B026 | HSCT | CH | RECTUM |
| **18** | B046 | HSCT | CI | RECTUM |
| **19** | B054 | HSCT | CIA | RECTUM |

| | | | | |
|---|---|---|---|---|
| **20** | B090 | HSCT | CH | SIGMOID COLON |
| **21** | B141 | HSCT | NA | SIGMOID COLON |
| **22** | B146 | HSCT | CI | SIGMOID COLON |
| **23** | B045 | HSCT | CH | TRANSVERSE COLON |
| **24** | B077 | HSCT | CIA | TRANSVERSE COLON |
| **25** | 1.H6 | Biotherapy | Colon | Colon |
| **26** | 1.H7 | Biotherapy | Colon | Colon |
| **27** | 1.H8 | Biotherapy | Colon | Colon |
| **28** | 1.H9 | Biotherapy | Ileorectal anastomosis | Ileorectal anastomosis |
| **29** | 1.I1 | Biotherapy | Ileorectal anastomosis | Ileorectal anastomosis |
| **30** | 1.I2 | Biotherapy | Ileorectal anastomosis | Ileorectal anastomosis |
| **31** | 1.A7 | Biotherapy | Ileum | Ileum |
| **32** | 1.A9 | Biotherapy | Ileum | Ileum |
| **33** | 1.B2 | Biotherapy | Ileum | Ileum |
| **34** | 1.B6 | Biotherapy | Ileum | Ileum |
| **35** | 1.B7 | Biotherapy | Ileum | Ileum |
| **36** | 1.B8 | Biotherapy | Ileum | Ileum |
| **37** | 1.E1 | Biotherapy | Ileum | Ileum |
| **38** | 1.E2 | Biotherapy | Ileum | Ileum |
| **39** | 1.E3 | Biotherapy | Ileum | Ileum |
| **40** | 2.A5 | Biotherapy | Ileum | Ileum |
| **41** | 2.A6 | Biotherapy | Ileum | Ileum |
| **42** | 2.A7 | Biotherapy | Ileum | Ileum |
| **43** | 2.B4 | Biotherapy | Ileum | Ileum |
| **44** | 2.B5 | Biotherapy | Ileum | Ileum |
| **45** | 2.B6 | Biotherapy | Ileum | Ileum |
| **46** | 2.D8 | Biotherapy | Ileum | Ileum |
| **47** | 2.D9 | Biotherapy | Ileum | Ileum |
| **48** | 2.E4 | Biotherapy | Ileum | Ileum |
| **49** | 2.E5 | Biotherapy | Ileum | Ileum |
| **50** | 2.E6 | Biotherapy | Ileum | Ileum |
| **51** | 1.A4 | Biotherapy | NA | NA |
| **52** | 1.D9 | Biotherapy | NA | NA |
| **53** | 1.E4 | Biotherapy | NA | NA |
| **54** | 1.E5 | Biotherapy | NA | NA |
| **55** | 1.E6 | Biotherapy | NA | NA |
| **56** | 1.H5 | Biotherapy | NA | NA |
| **57** | 2.I4 | Biotherapy | NA | NA |
| **58** | 2.I5 | Biotherapy | NA | NA |
| **59** | 2.I6 | Biotherapy | NA | NA |
| **60** | 3.C2 | Biotherapy | NA | NA |
| **61** | 3.C3 | Biotherapy | NA | NA |
| **62** | 1.G7 | Biotherapy | Rectum | Rectum |
| **63** | 1.G8 | Biotherapy | Rectum | Rectum |
| **64** | 2.B2 | Biotherapy | Rectum | Rectum |
| **65** | 2.B3 | Biotherapy | Rectum | Rectum |
| **66** | 2.D6 | Biotherapy | Rectum | Rectum |

| | | | | |
|---|---|---|---|---|
| **67** | 2.D7 | Biotherapy | Rectum | Rectum |
| **68** | 3.C7 | Biotherapy | Rectum | Rectum |
| **69** | 3.C8 | Biotherapy | Rectum | Rectum |
| **70** | 3.C9 | Biotherapy | Rectum | Rectum |
| **71** | 1.C8 | Biotherapy | Right Colon | Right Colon |
| **72** | 1.C9 | Biotherapy | Right Colon | Right Colon |
| **73** | 3.D7 | Biotherapy | Right Colon | Right Colon |
| **74** | 3.D8 | Biotherapy | Right Colon | Right Colon |
| **75** | 1.D6 | Biotherapy | Right colon or Rectum | Right colon or Rectum |
| **76** | 1.D7 | Biotherapy | Right colon or Rectum | Right colon or Rectum |
| **77** | 1.D8 | Biotherapy | Right colon or Rectum | Right colon or Rectum |
| **78** | 1.B9 | Biotherapy | Sigmoid | Sigmoid |
| **79** | 1.C1 | Biotherapy | Sigmoid | Sigmoid |
| **80** | 1.C4 | Biotherapy | Sigmoid | Sigmoid |
| **81** | 1.C5 | Biotherapy | Sigmoid | Sigmoid |
| **82** | 1.D1 | Biotherapy | Sigmoid | Sigmoid |
| **83** | 1.D2 | Biotherapy | Sigmoid | Sigmoid |
| **84** | 1.E8 | Biotherapy | Sigmoid | Sigmoid |
| **85** | 1.E9 | Biotherapy | Sigmoid | Sigmoid |
| **86** | 1.F1 | Biotherapy | Sigmoid | Sigmoid |
| **87** | 1.G5 | Biotherapy | Sigmoid | Sigmoid |
| **88** | 1.G6 | Biotherapy | Sigmoid | Sigmoid |
| **89** | 1.I3 | Biotherapy | Sigmoid | Sigmoid |
| **90** | 1.I4 | Biotherapy | Sigmoid | Sigmoid |
| **91** | 1.I5 | Biotherapy | Sigmoid | Sigmoid |
| **92** | 1.I6 | Biotherapy | Sigmoid | Sigmoid |
| **93** | 1.I7 | Biotherapy | Sigmoid | Sigmoid |
| **94** | 1.I8 | Biotherapy | Sigmoid | Sigmoid |
| **95** | 2.A3 | Biotherapy | Sigmoid | Sigmoid |
| **96** | 2.A4 | Biotherapy | Sigmoid | Sigmoid |
| **97** | 2.B7 | Biotherapy | Sigmoid | Sigmoid |
| **98** | 2.B8 | Biotherapy | Sigmoid | Sigmoid |
| **99** | 2.B9 | Biotherapy | Sigmoid | Sigmoid |
| **100** | 2.H2 | Biotherapy | Sigmoid | Sigmoid |
| **101** | 2.H3 | Biotherapy | Sigmoid | Sigmoid |
| **102** | 2.H4 | Biotherapy | Sigmoid | Sigmoid |
| **103** | 2.I7 | Biotherapy | Sigmoid | Sigmoid |
| **104** | 2.I8 | Biotherapy | Sigmoid | Sigmoid |
| **105** | 3.B3 | Biotherapy | Sigmoid | Sigmoid |
| **106** | 3.B4 | Biotherapy | Sigmoid | Sigmoid |
| **107** | 3.B5 | Biotherapy | Sigmoid | Sigmoid |
| **108** | 3.C4 | Biotherapy | Sigmoid | Sigmoid |
| **109** | 3.C5 | Biotherapy | Sigmoid | Sigmoid |
| **110** | 3.C6 | Biotherapy | Sigmoid | Sigmoid |
| **111** | 3.D1 | Biotherapy | Sigmoid | Sigmoid |
| **112** | 3.E1 | Biotherapy | Sigmoid | Sigmoid |

| | | | | |
|---|---|---|---|---|
| **113** | 2.I9 | Biotherapy | Transverse Colon | Transverse Colon |
| **114** | 3.A1 | Biotherapy | Transverse Colon | Transverse Colon |
| **115** | 3.A2 | Biotherapy | Transverse Colon | Transverse Colon |
| **116** | 046Ileum | Pediatric | 57,0 | Ileum |
| **117** | 046Sigma | Pediatric | 18,8 | Sigmoid |
| **118** | 008Colon | Pediatric | 4.22 | Sigmoid |
| **119** | 143Ileum | Pediatric | 52,2 | Ileum |
| **120** | 143Sigma | Pediatric | 17,1 | Sigmoid |
| **121** | B107 | HSCT | ControlCA | ASCENDING COLON |
| **122** | B109 | HSCT | ControlCA | ASCENDING COLON |
| **123** | B113 | HSCT | ControlCA | ASCENDING COLON |
| **124** | B115 | HSCT | ControlCA | ASCENDING COLON |
| **125** | B119 | HSCT | ControlCA | ASCENDING COLON |
| **126** | B120 | HSCT | ControlCA | ASCENDING COLON |
| **127** | B123 | HSCT | ControlCA | ASCENDING COLON |
| **128** | B126 | HSCT | ControlCA | ASCENDING COLON |
| **129** | B129 | HSCT | ControlCA | ASCENDING COLON |
| **130** | B132 | HSCT | ControlCA | ASCENDING COLON |
| **131** | B137 | HSCT | ControlCA | ASCENDING COLON |
| **132** | B140 | HSCT | ControlCA | ASCENDING COLON |
| **133** | B154 | HSCT | ControlCA | ASCENDING COLON |
| **134** | B157 | HSCT | ControlCA | ASCENDING COLON |
| **135** | B160 | HSCT | ControlCA | ASCENDING COLON |
| **136** | B168 | HSCT | ControlCA | ASCENDING COLON |
| **137** | B171 | HSCT | ControlCA | ASCENDING COLON |
| **138** | B122 | HSCT | ControlILI | ILEUM |
| **139** | B125 | HSCT | ControlILI | ILEUM |
| **140** | B128 | HSCT | ControlILI | ILEUM |
| **141** | B131 | HSCT | ControlILI | ILEUM |
| **142** | B134 | HSCT | ControlILI | ILEUM |
| **143** | B135 | HSCT | ControlILI | ILEUM |
| **144** | B138 | HSCT | ControlILI | ILEUM |
| **145** | B152 | HSCT | ControlILI | ILEUM |
| **146** | B155 | HSCT | ControlILI | ILEUM |
| **147** | B158 | HSCT | ControlILI | ILEUM |
| **148** | B169 | HSCT | ControlILI | ILEUM |
| **149** | B110 | HSCT | ControlSIG | SIGMOID COLON |
| **150** | B114 | HSCT | ControlSIG | SIGMOID COLON |
| **151** | B116 | HSCT | ControlSIG | SIGMOID COLON |
| **152** | B124 | HSCT | ControlSIG | SIGMOID COLON |
| **153** | B127 | HSCT | ControlSIG | SIGMOID COLON |
| **154** | B130 | HSCT | ControlSIG | SIGMOID COLON |
| **155** | B133 | HSCT | ControlSIG | SIGMOID COLON |
| **156** | B136 | HSCT | ControlSIG | SIGMOID COLON |
| **157** | B139 | HSCT | ControlSIG | SIGMOID COLON |
| **158** | B156 | HSCT | ControlSIG | SIGMOID COLON |
| **159** | B159 | HSCT | ControlSIG | SIGMOID COLON |

| | | | | |
|---|---|---|---|---|
| **160** | B167 | HSCT | ControlSIG | SIGMOID COLON |
| **161** | B170 | HSCT | ControlSIG | SIGMOID COLON |
| **162** | 010Colon | Pediatric | 7.04 | Sigmoid |
| **163** | 146Ileum | Pediatric | 71,3 | Ileum |
| **164** | 146Sigma | Pediatric | 17,6 | Sigmoid |
| **165** | B151 | HSCT | CIA | DESCENDING COLON |
| **166** | B150 | HSCT | IIA | ILEUM |
| **167** | B165 | HSCT | IIA | ILEUM |
| **168** | B164 | HSCT | CIA | SIGMOID COLON |
| **169** | 21952 | Pediatric | 1.85 | Ileum |
| **170** | 21982 | Pediatric | 1.88 | Ileum |
| **171** | 22715 | Pediatric | 1.86 | Ileum |
| **172** | 24266 | Pediatric | 1.6 | Ileum |
| **173** | 26181 | Pediatric | 1.87 | Ileum |
| **174** | 26919 | Pediatric | 1.91 | Ileum |
| **175** | 26951 | Pediatric | 1.69 | Ileum |
| **176** | 21967 | Pediatric | 1.87 | Sigma |
| **177** | 21997 | Pediatric | 1.64 | Sigma |
| **178** | 24296 | Pediatric | 1.7 | Sigma |
| **179** | 26197 | Pediatric | 2.01 | Sigma |
| **180** | 26213 | Pediatric | 1.98 | Sigma |
| **181** | 26229 | Pediatric | 1.86 | Sigma |
| **182** | 26935 | Pediatric | 1.68 | Sigma |
| **183** | 26967 | Pediatric | 1.74 | Sigma |
| **184** | 24250 | Pediatric | 1.85 | Sigma Transversum |
| **185** | B076 | HSCT | CIA | DESCENDING COLON |
| **186** | B100 | HSCT | CIA | DESCENDING COLON |
| **187** | B047 | HSCT | IH | ILEUM |
| **188** | B101 | HSCT | IH | ILEUM |
| **189** | B048 | HSCT | CIA | TRANSVERSE COLON |
| **190** | B014 | HSCT | CH | ASCENDING COLON |
| **191** | B033 | HSCT | CIA | ASCENDING COLON |
| **192** | B089 | HSCT | CIA | ASCENDING COLON |
| **193** | B103 | HSCT | CIA | ASCENDING COLON |
| **194** | B105 | HSCT | CIA | ASCENDING COLON |
| **195** | B073 | HSCT | CIA | DESCENDING COLON |
| **196** | B075 | HSCT | CII | DESCENDING COLON |
| **197** | B095 | HSCT | CH | DESCENDING COLON |
| **198** | B099 | HSCT | CIA | DESCENDING COLON |
| **199** | B013 | HSCT | IIA | ILEUM |
| **200** | B072 | HSCT | IH | ILEUM |
| **201** | B093 | HSCT | IIA | ILEUM |
| **202** | B094 | HSCT | IIA | ILEUM |
| **203** | B098 | HSCT | IH | ILEUM |
| **204** | B015 | HSCT | CII | RECTUM |
| **205** | B034 | HSCT | CH | RECTUM |
| **206** | B088 | HSCT | CH | SIGMOID COLON |

| | | | | |
|---|---|---|---|---|
| **207** | B102 | HSCT | CH | SIGMOID COLON |
| **208** | B104 | HSCT | CII | SIGMOID COLON |
| **209** | 1.D3 | Biotherapy | Ileum | Ileum |
| **210** | 1.D4 | Biotherapy | Ileum | Ileum |
| **211** | 1.D5 | Biotherapy | Ileum | Ileum |
| **212** | 2.C4 | Biotherapy | Ileum | Ileum |
| **213** | 2.C5 | Biotherapy | Ileum | Ileum |
| **214** | 2.C6 | Biotherapy | Ileum | Ileum |
| **215** | 2.G5 | Biotherapy | Ileum | Ileum |
| **216** | 2.G6 | Biotherapy | Ileum | Ileum |
| **217** | 2.G7 | Biotherapy | Ileum | Ileum |
| **218** | 3.B1 | Biotherapy | Ileum | Ileum |
| **219** | 3.B2 | Biotherapy | Ileum | Ileum |
| **220** | 3.D5 | Biotherapy | Ileum | Ileum |
| **221** | 3.D6 | Biotherapy | Ileum | Ileum |
| **222** | 3.E2 | Biotherapy | Ileum | Ileum |
| **223** | B050 | HSCT | III | ILEUM |
| **224** | B051 | HSCT | III | ILEUM |
| **225** | B040 | HSCT | CII | ASCENDING COLON |
| **226** | B042 | HSCT | CII | ASCENDING COLON |
| **227** | B163 | HSCT | CII | DESCENDING COLON |
| **228** | B038 | HSCT | IH | ILEUM |
| **229** | B043 | HSCT | IH | ILEUM |
| **230** | B161 | HSCT | IH | ILEUM |
| **231** | B039 | HSCT | CH | RECTUM |
| **232** | B041 | HSCT | CH | RECTUM |
| **233** | B162 | HSCT | CII | SIGMOID COLON |
| **234** | B006 | HSCT | CI | ASCENDING COLON |
| **235** | B008 | HSCT | CII | ASCENDING COLON |
| **236** | B011 | HSCT | CH | ASCENDING COLON |
| **237** | B017 | HSCT | CI | ASCENDING COLON |
| **238** | B018 | HSCT | CII | ASCENDING COLON |
| **239** | B021 | HSCT | CII | ASCENDING COLON |
| **240** | B023 | HSCT | CII | ASCENDING COLON |
| **241** | B028 | HSCT | CII | ASCENDING COLON |
| **242** | B030 | HSCT | CII | ASCENDING COLON |
| **243** | B036 | HSCT | CII | ASCENDING COLON |
| **244** | B063 | HSCT | CH | ASCENDING COLON |
| **245** | B143 | HSCT | CH | ASCENDING COLON |
| **246** | B069 | HSCT | CII | DESCENDING COLON |
| **247** | B070 | HSCT | CII | DESCENDING COLON |
| **248** | B081 | HSCT | CH | DESCENDING COLON |
| **249** | B084 | HSCT | CH | DESCENDING COLON |
| **250** | B085 | HSCT | CH | DESCENDING COLON |
| **251** | B005 | HSCT | II | ILEUM |
| **252** | B007 | HSCT | III | ILEUM |
| **253** | B012 | HSCT | III | ILEUM |

| | | | | |
|---|---|---|---|---|
| **254** | B016 | HSCT | II | ILEUM |
| **255** | B019 | HSCT | III | ILEUM |
| **256** | B020 | HSCT | III | ILEUM |
| **257** | B022 | HSCT | III | ILEUM |
| **258** | B027 | HSCT | IH | ILEUM |
| **259** | B031 | HSCT | IH | ILEUM |
| **260** | B035 | HSCT | IH | ILEUM |
| **261** | B055 | HSCT | III | ILEUM |
| **262** | B058 | HSCT | III | ILEUM |
| **263** | B059 | HSCT | III | ILEUM |
| **264** | B068 | HSCT | IH | ILEUM |
| **265** | B071 | HSCT | IH | ILEUM |
| **266** | B080 | HSCT | IIA | ILEUM |
| **267** | B083 | HSCT | III | ILEUM |
| **268** | B009 | HSCT | CH | RECTUM |
| **269** | B010 | HSCT | CII | RECTUM |
| **270** | B029 | HSCT | CH | RECTUM |
| **271** | B032 | HSCT | CH | RECTUM |
| **272** | B037 | HSCT | CH | RECTUM |
| **273** | B056 | HSCT | CII | RECTUM |
| **274** | B057 | HSCT | III | RECTUM |
| **275** | B060 | HSCT | CII | RECTUM |
| **276** | B062 | HSCT | CII | SIGMOID COLON |
| **277** | B064 | HSCT | CII | SIGMOID COLON |
| **278** | B142 | HSCT | CI | SIGMOID COLON |
| **279** | B144 | HSCT | CII | SIGMOID COLON |
| **280** | B079 | HSCT | CII | TRANSVERSE COLON |
| **281** | B082 | HSCT | CII | TRANSVERSE COLON |
| **282** | B086 | HSCT | CII | TRANSVERSE COLON |
| **283** | 2.H1 | Biotherapy | NA | NA |
| **284** | 1.C2 | Biotherapy | Rectum | Rectum |
| **285** | 1.C3 | Biotherapy | Rectum | Rectum |
| **286** | 1.E7 | Biotherapy | Rectum | Rectum |
| **287** | 1.F8 | Biotherapy | Rectum | Rectum |
| **288** | 1.F9 | Biotherapy | Rectum | Rectum |
| **289** | 1.G1 | Biotherapy | Rectum | Rectum |
| **290** | 1.G2 | Biotherapy | Rectum | Rectum |
| **291** | 1.G3 | Biotherapy | Rectum | Rectum |
| **292** | 1.G4 | Biotherapy | Rectum | Rectum |
| **293** | 2.C7 | Biotherapy | Rectum | Rectum |
| **294** | 2.C8 | Biotherapy | Rectum | Rectum |
| **295** | 2.D3 | Biotherapy | Rectum | Rectum |
| **296** | 2.D4 | Biotherapy | Rectum | Rectum |
| **297** | 2.D5 | Biotherapy | Rectum | Rectum |
| **298** | 2.E7 | Biotherapy | Rectum | Rectum |
| **299** | 2.E8 | Biotherapy | Rectum | Rectum |
| **300** | 3.A9 | Biotherapy | Rectum | Rectum |

| | | | | |
|---|---|---|---|---|
| **301** | 3.B8 | Biotherapy | Rectum | Rectum |
| **302** | 3.B9 | Biotherapy | Rectum | Rectum |
| **303** | 3.C1 | Biotherapy | Rectum | Rectum |
| **304** | 3.D9 | Biotherapy | Rectum | Rectum |
| **305** | 1.C6 | Biotherapy | Sigmoid | Sigmoid |
| **306** | 1.C7 | Biotherapy | Sigmoid | Sigmoid |
| **307** | 1.F2 | Biotherapy | Sigmoid | Sigmoid |
| **308** | 1.F3 | Biotherapy | Sigmoid | Sigmoid |
| **309** | 1.F4 | Biotherapy | Sigmoid | Sigmoid |
| **310** | 1.F5 | Biotherapy | Sigmoid | Sigmoid |
| **311** | 1.F6 | Biotherapy | Sigmoid | Sigmoid |
| **312** | 1.F7 | Biotherapy | Sigmoid | Sigmoid |
| **313** | 1.G9 | Biotherapy | Sigmoid | Sigmoid |
| **314** | 1.H1 | Biotherapy | Sigmoid | Sigmoid |
| **315** | 1.H2 | Biotherapy | Sigmoid | Sigmoid |
| **316** | 1.H3 | Biotherapy | Sigmoid | Sigmoid |
| **317** | 1.H4 | Biotherapy | Sigmoid | Sigmoid |
| **318** | 1.I9 | Biotherapy | Sigmoid | Sigmoid |
| **319** | 2.A1 | Biotherapy | Sigmoid | Sigmoid |
| **320** | 2.A2 | Biotherapy | Sigmoid | Sigmoid |
| **321** | 2.A8 | Biotherapy | Sigmoid | Sigmoid |
| **322** | 2.A9 | Biotherapy | Sigmoid | Sigmoid |
| **323** | 2.B1 | Biotherapy | Sigmoid | Sigmoid |
| **324** | 2.C1 | Biotherapy | Sigmoid | Sigmoid |
| **325** | 2.C2 | Biotherapy | Sigmoid | Sigmoid |
| **326** | 2.C3 | Biotherapy | Sigmoid | Sigmoid |
| **327** | 2.C9 | Biotherapy | Sigmoid | Sigmoid |
| **328** | 2.D1 | Biotherapy | Sigmoid | Sigmoid |
| **329** | 2.D2 | Biotherapy | Sigmoid | Sigmoid |
| **330** | 2.E1 | Biotherapy | Sigmoid | Sigmoid |
| **331** | 2.E2 | Biotherapy | Sigmoid | Sigmoid |
| **332** | 2.E3 | Biotherapy | Sigmoid | Sigmoid |
| **333** | 2.E9 | Biotherapy | Sigmoid | Sigmoid |
| **334** | 2.F1 | Biotherapy | Sigmoid | Sigmoid |
| **335** | 2.F2 | Biotherapy | Sigmoid | Sigmoid |
| **336** | 2.F3 | Biotherapy | Sigmoid | Sigmoid |
| **337** | 2.F4 | Biotherapy | Sigmoid | Sigmoid |
| **338** | 2.F5 | Biotherapy | Sigmoid | Sigmoid |
| **339** | 2.F6 | Biotherapy | Sigmoid | Sigmoid |
| **340** | 2.F7 | Biotherapy | Sigmoid | Sigmoid |
| **341** | 2.F8 | Biotherapy | Sigmoid | Sigmoid |
| **342** | 2.F9 | Biotherapy | Sigmoid | Sigmoid |
| **343** | 2.G1 | Biotherapy | Sigmoid | Sigmoid |
| **344** | 2.G2 | Biotherapy | Sigmoid | Sigmoid |
| **345** | 2.G3 | Biotherapy | Sigmoid | Sigmoid |
| **346** | 2.G4 | Biotherapy | Sigmoid | Sigmoid |
| **347** | 2.G8 | Biotherapy | Sigmoid | Sigmoid |

| | | | | |
|---|---|---|---|---|
| **348** | 2.G9 | Biotherapy | Sigmoid | Sigmoid |
| **349** | 2.H5 | Biotherapy | Sigmoid | Sigmoid |
| **350** | 2.H6 | Biotherapy | Sigmoid | Sigmoid |
| **351** | 2.H7 | Biotherapy | Sigmoid | Sigmoid |
| **352** | 2.H8 | Biotherapy | Sigmoid | Sigmoid |
| **353** | 2.H9 | Biotherapy | Sigmoid | Sigmoid |
| **354** | 2.I1 | Biotherapy | Sigmoid | Sigmoid |
| **355** | 2.I2 | Biotherapy | Sigmoid | Sigmoid |
| **356** | 2.I3 | Biotherapy | Sigmoid | Sigmoid |
| **357** | 3.A3 | Biotherapy | Sigmoid | Sigmoid |
| **358** | 3.A4 | Biotherapy | Sigmoid | Sigmoid |
| **359** | 3.A5 | Biotherapy | Sigmoid | Sigmoid |
| **360** | 3.A6 | Biotherapy | Sigmoid | Sigmoid |
| **361** | 3.A7 | Biotherapy | Sigmoid | Sigmoid |
| **362** | 3.A8 | Biotherapy | Sigmoid | Sigmoid |
| **363** | 3.B6 | Biotherapy | Sigmoid | Sigmoid |
| **364** | 3.B7 | Biotherapy | Sigmoid | Sigmoid |
| **365** | 3.D2 | Biotherapy | Sigmoid | Sigmoid |
| **366** | 3.D3 | Biotherapy | Sigmoid | Sigmoid |
| **367** | 3.D4 | Biotherapy | Sigmoid | Sigmoid |
| **368** | 048Ileum | Pediatric | 138,3 | Ileum |
| **369** | 049Ileum | Pediatric | 153,5 | Ileum |
| **370** | 066Ileum | Pediatric | 17,9 | Ileum |
| **371** | 147Ileum | Pediatric | 62,8 | Ileum |
| **372** | 149Ileum | Pediatric | 60,0 | Ileum |
| **373** | 150Ileum | Pediatric | 86,8 | Ileum |
| **374** | 140Ileum | Pediatric | 203,1 | Ileum |
| **375** | 144Ileum | Pediatric | 10,1 | Ileum |
| **376** | 023Ileum | Pediatric | 3.81 | Ileum |
| **377** | 048Sigma | Pediatric | 13,6 | Sigmoid |
| **378** | 049Sigma | Pediatric | 19,4 | Sigmoid |
| **379** | 050Sigma | Pediatric | 116,4 | Sigmoid |
| **380** | 149Sigma | Pediatric | 82,6 | Sigmoid |
| **381** | 066Sigma | Pediatric | 67,3 | Sigmoid |
| **382** | 140Sigma | Pediatric | 14,5 | Sigmoid |
| **383** | 144Sigma | Pediatric | 124,4 | Sigmoid |
| **384** | 023Colon | Pediatric | 1.46 | Sigmoid |
| **385** | 22013 | Pediatric | 1.86 | Ileum |
| **386** | 22043 | Pediatric | 1.77 | Ileum |
| **387** | 22073 | Pediatric | 1.85 | Ileum |
| **388** | 22104 | Pediatric | 1.83 | Ileum |
| **389** | 25026 | Pediatric | 1.84 | Ileum |
| **390** | 24998 | Pediatric | 1.91 | Ileum |
| **391** | 24859 | Pediatric | 1.89 | Ileum |
| **392** | 24891 | Pediatric | 1.84 | Ileum |

| | | | | |
|---|---|---|---|---|
| **393** | 24928 | Pediatric | 1.81 | Ileum |
| **394** | 24960 | Pediatric | 1.89 | Ileum |
| **395** | 26390 | Pediatric | 1.91 | Ileum |
| **396** | 26608 | Pediatric | 1.86 | Ileum |
| **397** | 26447 | Pediatric | 1.92 | Ileum |
| **398** | 22028 | Pediatric | 1.8 | Sigma |
| **399** | 22058 | Pediatric | 2.23 | Sigma |
| **400** | 22089 | Pediatric | 1.8 | Sigma |
| **401** | 25040 | Pediatric | 1.87 | Sigma |
| **402** | 25012 | Pediatric | 1.41 | Sigma |
| **403** | 24875 | Pediatric | 1.86 | Sigma |
| **404** | 24907 | Pediatric | 1.84 | Sigma |
| **405** | 24944 | Pediatric | 2.02 | Sigma |
| **406** | 24976 | Pediatric | 1.91 | Sigma |
| **407** | 26406 | Pediatric | 1.6 | Sigma |
| **408** | 26624 | Pediatric | 1.89 | Sigma |
| **409** | 26429 | Pediatric | 1.84 | Sigma |
| **410** | 26463 | Pediatric | 1.87 | Sigma |
| **411** | 1.B3 | Biotherapy | Ileum | Ileum |
| **412** | 1.B4 | Biotherapy | Ileum | Ileum |
| **413** | 1.B5 | Biotherapy | Ileum | Ileum |
| **414** | B049 | HSCT | | Colon |
| **415** | B065 | HSCT | | Colon |
| **416** | B173 | HSCT | 1.81 | Colon |
| **417** | B176 | HSCT | 1.84 | Colon |
| **418** | B177 | HSCT | 1.86 | Colon |
| **419** | B178 | HSCT | 1.83 | Colon |
| **420** | B179 | HSCT | 1.81 | Colon |
| **421** | B181 | HSCT | 1.82 | Colon |
| **422** | B182 | HSCT | 1.82 | Colon |
| **423** | B183 | HSCT | 1.78 | Colon |
| **424** | B172 | HSCT | 1.86 | Ileum |
| **425** | B174 | HSCT | 1.88 | Ileum |
| **426** | B175 | HSCT | 1.77 | Ileum |
| **427** | B180 | HSCT | 1.75 | Ileum |
| **428** | B149 | HSCT | NA | Ileum |
| **429** | B184 | HSCT | 1.89 | Ileum |
| **430** | B185 | HSCT | 1.9 | Ileum |

**Supplementary Table S3:** Minimal bacterial consortium (MIBAC) composition

| *Species* | Original strain designation | DSM no. |
|---|---|---|
| ***Clostridium ramosum*** | **SRB509'-5-F-B** | **29357** |
| ***Paraclostridium bifermentans*** | **G7K1R3-PYG-90** | **29423** |
| ***Enterococcus hirae*** | **SB** | **28619** |
| ***Enterorhabdus mucosicola*** | **Mt1B8** | **19490** |
| ***Escherichia coli*** | **Mt1B1** | **28618** |
| ***Lactobacillus murinus*** | **M-6244-3B** | **28683** |
| ***Parabacteroides goldsteinii*** | **BS-C3-2** | **29187** |
